# Evaluating functional brain organization in individuals and identifying contributions to network overlap

**DOI:** 10.1101/2023.09.21.558809

**Authors:** Janine D. Bijsterbosch, Seyedeh-Rezvan Farahibozorg, Matthew F. Glasser, David Van Essen, Lawrence H. Snyder, Mark W. Woolrich, Stephen M. Smith

## Abstract

Individual differences in the spatial organization of resting state networks have received increased attention in recent years. Measures of individual-specific spatial organization of brain networks and overlapping network organization have been linked to important behavioral and clinical traits and are therefore potential biomarker targets for personalized psychiatry approaches. To better understand individual-specific spatial brain organization, this paper addressed three key goals. First, we determined whether it is possible to reliably estimate weighted (non-binarized) resting state network maps using data from only a single individual, while also maintaining maximum spatial correspondence across individuals. Second, we determined the degree of spatial overlap between distinct networks, using test-retest and twin data. Third, we systematically tested multiple hypotheses (spatial mixing, temporal switching, and coupling) as candidate explanations for why networks overlap spatially. To estimate weighted network organization, we adopt the Probabilistic Functional Modes (PROFUMO) algorithm, which implements a Bayesian framework with hemodynamic and connectivity priors to supplement optimization for spatial sparsity/independence. Our findings showed that replicable individual-specific estimates of weighted resting state networks can be derived using high quality fMRI data within individual subjects. Network organization estimates using only data from each individual subject closely resembled group-informed network estimates (which was not explicitly modeled in our individual-specific analyses), suggesting that cross-subject correspondence was largely maintained. Furthermore, our results confirmed the presence of spatial overlap in network organization, which was replicable across sessions within individuals and in monozygotic twin pairs. Intriguingly, our findings provide evidence that network overlap is indicative of linear additive coupling. These results suggest that regions of network overlap concurrently process information from all contributing networks, potentially pointing to the role of overlapping network organization in the integration of information across multiple brain systems.

## 1. Introduction

Recent studies have revealed substantial inter-individual variability in the spatial organization of the brain as measured with resting state functional MRI (rfMRI) (Braga & Buckner, 2017; Glasser et al., 2016; Gordon, Laumann, Adeyemo, & Petersen, 2017; Gordon, Laumann, Adeyemo, Gilmore, et al., 2017; Gordon, Laumann, Gilmore, et al., 2017; Laumann et al., 2015; D. Wang et al., 2015). Importantly, such inter-individual spatial variability in functional brain organization is strongly associated with behavioral traits (Bijsterbosch et al., 2018; Kong et al., 2019). The overarching objective of this paper is to characterize weighted (i.e., non-binarized) spatial organization of resting state networks within individuals, with a specific focus on gaining insights into spatially overlapping network organization.

Identifying network organization at the individual level rather than using group information raises multiple challenges. First, individual estimates of network organization are noisier than group estimates, partly because group estimates benefit from collating data across participants. This challenge can be partially addressed by obtaining large amounts of data from each individual (precision functional mapping approach (Gratton et al., 2020)), but such extensive data acquisition may not be feasible in all participants and settings. Second, although a purely individual-specific set of network maps represents the most accurate and unbiased estimate of the individual’s brain organization (Gordon, Laumann, Gilmore, et al., 2017), it may lack network correspondence across individuals. Assuming the presence of cross-participant commonalities in their network structure, such group correspondence is valuable for network labeling and interpretability, and essential for group-level and between-subject analytical comparisons. Group-based estimates applied to individuals have built-in correspondence, but these individual estimates may be biased towards the group estimate (Bijsterbosch et al., 2019). Probabilistic Functional Modes (PROFUMO) is a Hierarchical Bayesian algorithm developed to try to optimize this trade-off by using group-level priors to achieve correspondence, whilst optimizing individual-specific estimates to maximize the accuracy of individual network maps (Harrison et al., 2015). Although PROFUMO has been successfully applied in group data (Bijsterbosch et al., 2018; Farahibozorg et al., 2021; Harrison et al., 2020), an open question is whether it can be robustly applied to data from only a single individual without sacrificing correspondence, which is of particular interest in the context of personalized psychiatry and translational work in non-human primates and other animal models. The first goal of this paper was to determine whether PROFUMO can reliably estimate weighted (i.e., non-binarized) resting state networks using only data from a single subject, while also achieving substantial “non-enforced” correspondence across individuals (i.e., without an explicit hierarchical model for correspondence).

Spatial overlap between rfMRI networks beyond classical ‘hub regions’ has been observed across a variety of analytical brain representations (Lee et al., 2016; Lin et al., 2018; Najafi et al., 2016), and has been linked to behavioral traits (Bijsterbosch et al., 2019). PROFUMO accurately estimates spatial overlap in rfMRI network organization (Bijsterbosch et al., 2019), which is a key advantage compared to approaches that aim for a ‘hard’ binarized parcellation or approaches that enforce spatial independence between networks (Bijsterbosch et al., 2020). Despite broad interest in ‘hub’ regions within the graph theory domain that typically adopts hard parcellations (Bertolero et al., 2018; Buckner et al., 2009; Warren et al., 2014), a detailed spatial investigation into individual-specific overlap between weighted estimates of network organization is lacking. Studying the overlapping properties of individualized brain networks is of interest because network overlap may be a sensitive marker for use in personalized psychiatry settings (Gratton et al., 2020; Insel, 2014; Williams, 2016) given prior evidence of behavioral relevance (Bijsterbosch et al., 2019), provided that it can be robustly and reliably detected in individuals. Furthermore, this work contributes to the broader literature on precision functional mapping (Gordon, Laumann, Gilmore, et al., 2017; Gratton et al., 2018; Poldrack, 2017; Poldrack et al., 2015). The second goal of this paper was to characterize network overlap among weighted individual-specific resting state networks estimated using PROFUMO.

Each type of resting state fMRI analysis provides a different low-dimensional representation of the dataset (Bijsterbosch et al., 2020). Although optimized to best fit the data (Bijsterbosch et al., 2021), these brain representations are necessarily lossy given the intrinsic goal of dimensionality reduction. In the case of PROFUMO, a set of stationary large-scale modes of brain organization are derived that collapse fine grained spatial structure and simplify temporally dynamic processes. As such, there are multiple potential mechanisms that may give rise to spatially overlapping network organization as observed between network maps estimated with PROFUMO. First, it is possible that a brain region in which two networks appear to overlap may in fact be a spatially heterogeneous mixture of cortical patches that are individually part of either network 1 or network 2, implying no real functional “link” between the two networks as a result of the overlap (Fig. 1A). Such a spatially heterogeneous overlap region may, for example, result from network interdigitation (Braga et al., 2019; Braga & Buckner, 2017), or regional gradients (Blazquez Freches et al., 2020; Haak et al., 2018). Second, the region of network overlap may dynamically switch its network allegiance over time to be part of either network 1 or network 2 at any given moment (Hutchison et al., 2013) (Fig. 1B). Such dynamic switching would appear as a spatially overlapping network structure given the stationary (time-averaging) nature of the PROFUMO model. Third, network overlap might indicate that signals from network 1 and network 2 are jointly processed and “deeply functionally integrated” within regions of network overlap (Fig. 1C). This third hypothesis is perhaps the most intriguing as it may indicate information coupling involving a specific functional role of overlap regions, and contributing to between-network communication. The third goal of this paper was to systematically compare these spatial mixture, dynamic switching, and coupling hypotheses of network overlap.

**Figure 1:**
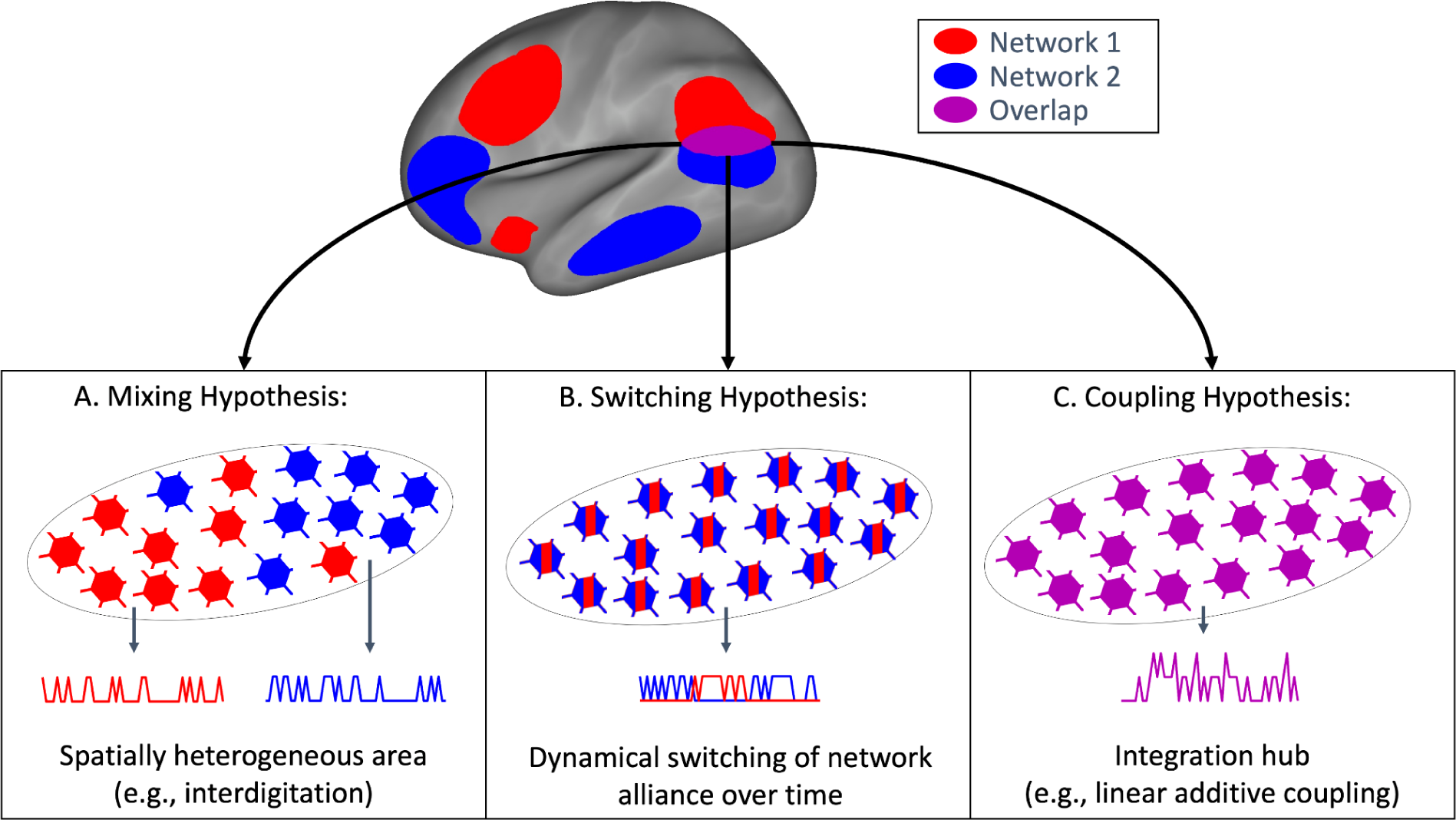
Graphical summary of the three hypotheses of network overlap. A) The spatial mixing hypothesis suggests that individual vertices (illustrated as neurons) within a region of network overlap may be part of either Network 1 (red) or Network 2 (blue). B) The dynamic switching hypothesis suggests that a region of network overlap may be spatially homogeneous but may switch network allegiance between Network 1 (red) and Network 2 (blue) over time. C) The coupling hypothesis suggests that a region of network overlap may integrate information from both Networks 1 and 2.

In this paper we leveraged a unique subset of the Human Connectome Project Young Adult data (Van Essen et al., 2013), by focusing on individuals who underwent three complete resting state sessions (3T, 3T re-scan, and 7T) for a total of approximately 3 hours of rfMRI data per individual. The resulting sample of N=20 that met this criteria further included 8 monozygotic twin pairs, providing a rich cohort to investigate individual-specific weighted network organization. Our focus on a small sample of densely sampled individuals was informed by the interest in individual-specific network organization. Although no brain-behavior associations were feasible given the small sample (Marek et al., 2022), prior work has extensively studied individual differences in PROFUMO including behavioral associations with spatial organization (Bijsterbosch et al., 2018), heritability (Harrison et al., 2020), and network variants as a function of dimensionality (Farahibozorg et al., 2021). The results of this work support the application of PROFUMO weighted networks in individual participants, which paves the way for future applications in a personalized psychiatry framework. Furthermore, our findings suggest a linearly additive coupling mechanism underlying spatially overlapping network organization, which supports the hypothesis that regions of network overlap play an important functional role in terms of cross-network coupling (Gordon et al., 2018).

## 2. Methods

### 2.1 Dataset

We used high quality data from the Human Connectome Project (HCP) (Van Essen et al., 2013), focusing on N=20 individuals (including 8 monozygotic twin pairs) who underwent a complete set of four 3T, four 7T and four retest 3T runs, thereby accumulating approximately three hours (13,200 timepoints) of rfMRI data across 12 scans per individual. This HCP-YA sub-sample was 80% female with a mean age of 30.1 years (standard deviation = 3.84; range 22-34). Briefly, the 3T rfMRI data were acquired at 2mm isotropic voxel size using a multiband factor of 8, a TR of 0.72 seconds, and a TE of 33 ms; the 7T rfMRI were acquired at 1.6mm isotropic voxel size, a multiband acceleration of 5, in-plane acceleration 2, a TR of 1.0 seconds, and a TE of 22.2ms (see further details in (Smith et al., 2013; T Vu et al., 2017; Van Essen et al., 2013)). Data were preprocessed using the HCP minimal processing pipelines (Glasser et al., 2013). All 3T and 7T data were analyzed in the Connectivity Informatics Technology Initiative (CIFTI) format, which consists of 91,282 grayordinates with approximately 2-mm spatial resolution (i.e., 7T data were downsampled to match the 3T spatial resolution). ICA-FIX was then applied to remove structured noise (Griffanti et al., 2014; Salimi-Khorshidi et al., 2014), and data were aligned using multimodal surface matching (MSM-All; (Robinson et al., 2014)) to align areal features (myelin and RSNs).

### 2.2 PROFUMO estimation

PROFUMO is a matrix factorization approach for the estimation of resting state networks that adopts spatial prior, temporal priors, and a noise model in a hierarchical Bayesian model (see (Harrison et al., 2015, 2020) and Supplementary Tables 1 and 2). PROFUMO was applied in several distinct ways:

1. Classic group-PROFUMO was performed in which data from all 12 runs across all 20 participants were used and modeled hierarchically according to the levels of subjects and (beneath that) runs. Notably, this version of PROFUMO is recommended for studies that include only one or a small number of sessions per individual.
2. Single-subject PROFUMO was performed independently for each of the 20 participants, using all 12 runs for each, those 12 being considered as separate ‘subjects’ in the estimation of PROFUMO’s Gaussian mixture model (Harrison et al., 2020).
3. Classic group-PROFUMO was performed similar to case (#1), using 12 individual runs from 12 separate participants. This was in order to obtain a group-level estimation of modes using the same amount of data available for the separate subject runs (case (#2), i.e., matching the effective signal to noise ratio; SNR).
4. Test-retest single-subject PROFUMO was performed independently for each of the 20 participants and independently using two sets of 6 runs each (split evenly across 3T, 7T, and retest data; and always including a pair of opposite phase-encode directions).
5. Single-subject PROFUMO similar to case (#2) was performed stepwise on cut-down versions of the 12 runs that included a progressively increasing number of timepoints (in increments of 1/12th of the run timepoints) to determine how much data are needed to obtain reliable network estimates. This approach was taken to ensure that all PROFUMO runs included equal contributions from 3T, retest, and 7T data across all phase-encode directions.

All of the PROFUMO runs were performed at a dimensionality of 20 to focus on the spatial organization of large scale resting state networks. Furthermore, classic group-PROFUMO (#1) was repeated at dimensionalities 30, 40, and 50 to determine the stability of the selected networks across PROFUMO decomposition dimensionalities. Beyond the key parameter of dimensionality (number of networks), all PROFUMO parameters were set to the default. A summary of all PROFUMO parameters, hyperparameters, and hyperpriors can be seen in Supplementary Tables S1 and S2. Beyond these PROFUMO parameters, additional key measures of interest for this paper include the similarity between group and individual network estimates indicative of correspondence and the test-retest reliability of network spatial maps representative of stable and reliable network estimates (described further below in sections 2.3 and 2.4).

### 2.3 PROFUMO mode selection

The Hungarian algorithm (a.k.a. ‘munkres’ algorithm) was used to reorder PROFUMO modes (i.e., “networks”) for each of the single-subject runs (#2 above), the split 1 and split 2 single-subject runs (#4 above), and the 12-run group run (#3 above) to best-match the mode order obtained from the full group results. Briefly, the Hungarian algorithm solves the assignment problem by permuting rows to minimize the trace of the permuted cost matrix (Kuhn, 1955; Munkres, 1957). For each of the 20 modes we estimated the test-retest correlation per subject as the Pearson correlation across all 91,282 CIFTI grayordinates between split 1 and split 2 single-subject runs. For each of the 20 modes we also estimated the subject-group correlation per subject as the Pearson correlation across all 91,282 vertices between the map from the subject run (#2 above) and the subject-specific estimated map from the group run (#1 above). This ‘group-individual run’ measure addresses the question of correspondence across individuals because it compares classic group PROFUMO modes (which benefit from explicit correspondence through the group prior in the hierarchical Bayesian algorithm) to single-subject PROFUMO modes (which do not involve any group information). Modes that achieved both a median (across subjects) test-retest correlation of 0.7 or greater and a median (across subjects) group-individual correlation of 0.7 or greater were used in subsequent analyses. Out of the 20 PROFUMO modes, 12 modes met these requirements. Within these 12 modes, individual participants’ missing modes were defined as modes with both a test-retest correlation lower than 0.2 and a subject-group correlation lower than 0.2 (both estimated at the subject level). Missing modes were ignored in subsequent analyses. For naming purposes, modes were spatially mapped onto the Yeo-7 parcellation (Yeo et al., 2011) and we followed the network naming taxonomy suggested by (Uddin et al., 2019).

### 2.4 PROFUMO mode stability

We assessed the stability of spatial maps in several ways. As described in section 2.3, the first two measures of mode stability were within-participant *test-retest reliability* between splits 1 and 2, and *within-subject similarity (group-individual run)* correlations between the subject run (#2 above) and the subject estimates obtained in the group run (#1 above). We also tested *twin similarity (individual runs)* and *twin similarity (group run)* by correlating each PROFUMO mode within each monozygotic twin pair using modes from separate subject runs (#2 above) or subject-specific estimates from the group run (#1 above), respectively. For comparison, we also investigated *between-subject similarity (individual runs)* and *between-subject similarity (group run),* reflecting the same across-subject correlations for all possible non-twin pairs of participants. All mode stability measures were performed on each spatial map (correlation across 91,282 vertices), on the temporal connectivity matrix (correlation across the 66 edges in the lower triangle, where edges represent the partial correlation between mode timeseries), and on the spatial overlap matrix (correlation across the 66 edges in the lower triangle, where edges represent the correlation between mode maps; see section 2.5 and (Bijsterbosch et al., 2019)).

### 2.5 Spatial overlap measures

To quantify spatial overlap across all twelve modes, we generated a spatial overlap matrix by estimating the Pearson’s correlation coefficients across grayordinates between all possible pairs across the 12 network maps, as developed in (Bijsterbosch et al., 2019). Correlation coefficients were z-transformed prior to averaging across individuals. Stability of individual specific spatial overlap matrices was calculated as described in section 2.4. To estimate spatial maps of network overlap in individual participants, each mode was binarized using a threshold of 1, and the number of overlapping modes were counted at each vertex. A threshold of 1 represents the 96.5^th^ percentile across all map weights and therefore offers a relatively conservative estimate of spatial overlap that is driven only by vertices with strong network contributions.

### 2.6 Focusing on 2-Network overlap

To systematically compare our hypotheses regarding network overlap, we focused on spatial overlap between pairs of networks. For each individual and each possible network pair (12!=66), we identified vertices uniquely associated with network 1 as those vertices with a weight of 1 or greater for network 1, a weight of less than 0.1 for network 2, and a summed weight across all other 10 modes of less than 0.1 (Fig. 2B). Negative map weights were not included in the estimation of network overlap, because the inclusion of negative vertices would dilute averaged timeseries by canceling out positive vertices. A similar procedure was used to identify vertices uniquely associated with network 2. To locate the spatial overlap region, we identified vertices having a weight of 1 or greater for both network 1 and network 2 and a summed weight across all other 10 modes of less than 0.1. This procedure was performed using mode maps estimated from the individual subject PROFUMO runs (#2 above). PROFUMO spatial maps are calculated by vertex-wise multiplying the probability (0-1) by the estimated mean (derived from Gaussian mixture model). Resulting spatial map values in our data ranged from −3.6 to 7.1, and the applied threshold of 1 represents the 96.4th percentile. Hence, thresholds were applied to the PROFUMO spatial maps as estimated by the PROFUMO algorithm without further transforms. As such, we focused specifically on two-mode overlap vertices, by removing vertices with significant contributions of additional modes (Fig. 2A). We focus on two-network overlap because it offers a tightly controlled test-bed for the hypothesis testing element of our work. Future work may expand into more complex overlapping network organization. Overlap regions were defined based on the data-driven approach described above without any explicit exclusions, and the resulting overlap patterns (see Supplementary Figure S1) comprehensively cover all areas of potential interest.

**Figure 2:**
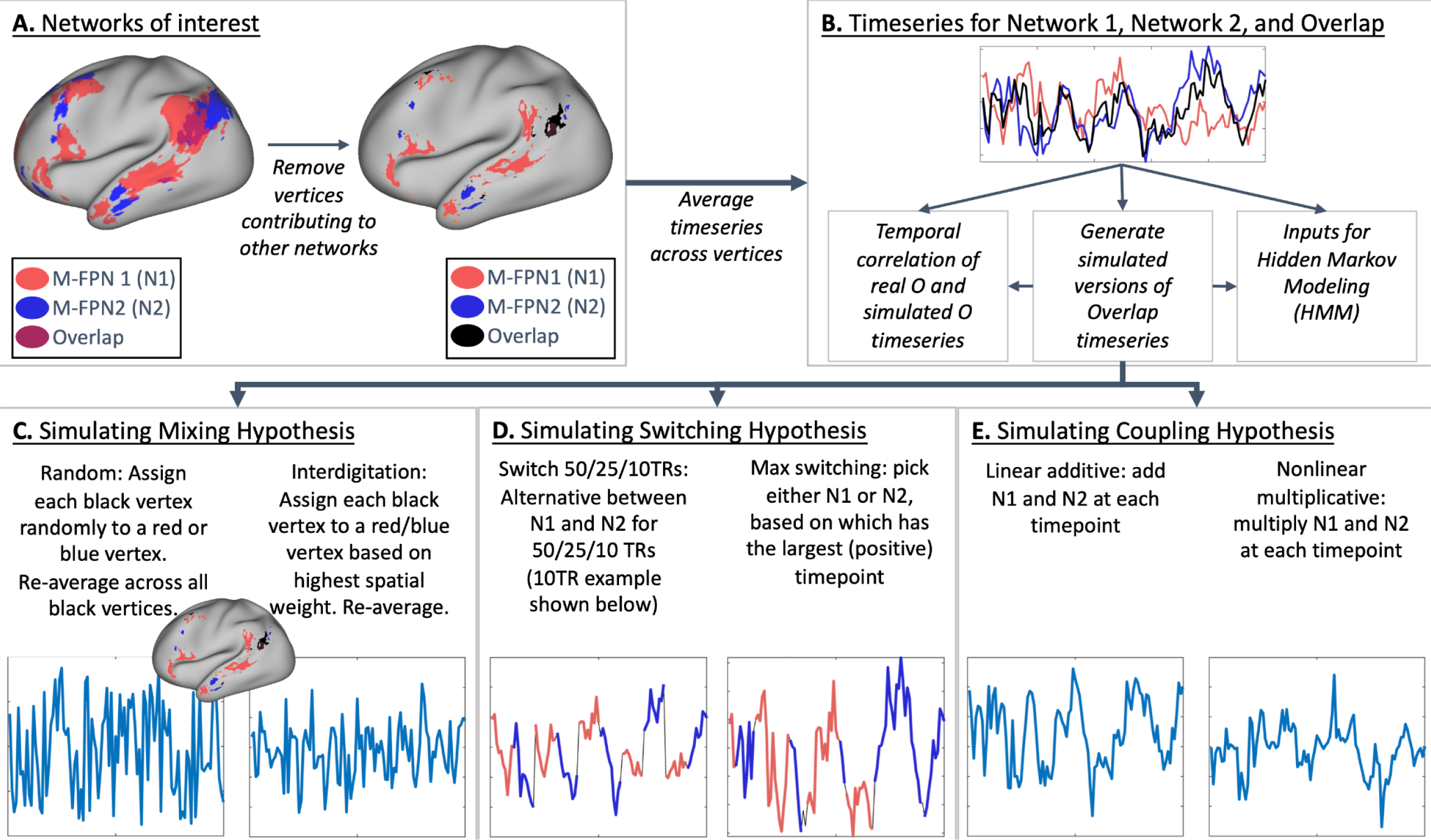
Graphical summary of overlap hypothesis testing methodology. A) Two spatially overlapping networks are selected and vertices are assigned to either network 1, network 2, or network Overlap. Vertices contributing to additional networks are excluded. B) Average timeseries are extracted per subject, per run for Network 1, Network 2, and Overlap. C) Mixing hypothesis-based semi-simulated (i.e., using actual timeseries from non-overlap vertices) versions of the Overlap timeseries are generated based on random and interdigitation-based mixtures of Network 1 and Network 2 vertices. D) Switching hypothesis-based semi-simulated versions of the Overlap timeseries are generated based on concatenated segments from Network 1 and 2 timeseries. E) Coupling hypothesis-based semi-simulated versions of the Overlap timeseries are generated based on additive or multiplicative combinations of Network 1 and 2 timeseries.

The number of network pairs for further investigation was reduced from 66 (all possible pairs) by selecting only those networks pairs in which the spatial overlap regions contained at least 25 vertices for at least half of the participants (n>=10). This network pair selection was performed to focus on pairs with robust and replicable overlap, and to reduce the computational demands for subsequent analyses. Out of the 66 possible network pairs, 20 pairs were selected for further analysis (see Supplementary Figure S1).

For each of the 20 network pairs, three summary timeseries were calculated (Fig. 2B) by averaging across: all vertices uniquely associated with network 1 (“N1”), all vertices uniquely associated with network 2 (“N2”), and all vertices in the overlap region (“O”). Each timeseries was standardized to a mean of zero and a standard deviation of 1 within each of the 12 runs. Any participants who did not have any vertices in the overlap region based on these criteria and were excluded from subsequent analyses.

### 2.7 Generating semi-simulated data for the overlap region

Multiple mechanistic hypotheses might in principle explain the occurrence of spatial overlap in stationary maps estimated using PROFUMO, and it is currently unknown what drives the observed (apparent) spatial overlap. To address this issue, we generated semi-simulated versions of overlap timeseries to test different hypotheses as described below. The semi-simulated version of overlap timeseries were compared to the original overlap timeseries using direct timeseries correlation, frequency characteristics (see section 2.8 for further details), and state means estimated with Hidden Markov Modeling (HMM; see section 2.9 for further details).

#### 2.7.1 Network switching hypothesis

One hypothesis is that the overlap region may dynamically switch network alliance between networks 1 and 2 over time within a scanning run (Fig. 1B), with the constraint that one and only one network is “active” at any given time (this is also a common constraint in the HMM modeling). Such dynamic switching would appear as overlap when using stationary methods such as PROFUMO, which effectively average across time. To test this hypothesis, we semi-simulated four versions of the overlap timeseries using the real N1 and N2 timeseries for each participant and run:

- <u>Switch 50 TRs (Fig. 2D)</u> starts with the first 50 timepoints from N1, then contains the second 50 timepoints from N2, then the third 50 timepoints from N1 and so on. Notably, potential phase shifts between original and semi-simulated data (i.e., mismatches in the order between N1 and N2) may impact the correlation between the original and semi-simulated versions of the timeseries. However, such phase discrepancies would not impact our second comparison measure based on HMM state means, because these are estimated independently of state timecourses.
- <u>Switch 25 TRs (Fig. 2D)</u> is the same as above, but switching between N1 and N2 every 25 timepoints.
- <u>Switch 10 TRs (Fig. 2D)</u> is the same as above, but switching between N1 and N2 every 10 timepoints.
- <u>Max switching (Fig. 2D)</u> assigns each timepoint in the semi-simulated overlap timeseries as the maximum from either N1 or N2 based on whichever datapoint (in real data timeseries N1 and N2) is higher for a given TR.

#### 2.7.2 Coupling hypothesis

Another hypothesis is that the overlap region is integrating data from both network 1 and network 2 at each TR (Fig. 1C). To test the coupling hypothesis, we semi-simulated two versions of the overlap timeseries using the real N1 and N2 timeseries for each participant and run:

- <u>Linear additive coupling (Fig. 2E)</u> is the sum of N1 and N2 within each TR.
- <u>Nonlinear multiplicative coupling (Fig. 2E)</u> takes the element-wise product between N1 and N2 after setting N1 and N2 to zero min respectively.

#### 2.7.3 Spatial mixture hypothesis

A final hypothesis is that the overlap region is a spatial mixture of vertices linked to network 1 and network 2 (Fig. 1A), akin to the concept of network interdigitation (Braga & Buckner, 2017) or within-region gradients (Haak et al., 2018). To test this hypothesis, we semi-simulated two versions of the overlap timeseries using the real vertex timeseries uniquely associated with either network 1 or network 2. In contrast to the other semi-simulated versions of the overlap timeseries described above, this method does not use the mean N1 and N2 timeseries and instead repeats the averaging across vertices.

- <u>Spatial random mixture (Fig. 2C)</u> assigns half of the vertices in the overlap region to a randomly chosen vertex out of those uniquely associated with network 1 (with replacement) and assigns the other half of the vertices in the overlap region to a randomly chosen vertex out of those uniquely associated with network 2 (with replacement). The semi-simulated overlap timeseries is then averaged across all vertices and standardized as above.
- <u>Spatial interdigitation (Fig. 2C)</u> assigns each vertex in the overlap region based on whether the spatial weight for that vertex was higher for network 1 or for network 2. If the spatial weight for M-FPN 1 is higher, the timeseries of the vertex uniquely associated with network 1 with the grayordinate that is closest in vectorized indexing is assigned to that overlap vertex (with replacement). The semi-simulated overlap timeseries is then averaged across all overlap vertices and standardized as above. We refer to this option as ‘spatial interdigitation’ because the PROFUMO spatial weights reflect spatially contiguous areas (see supplementary figure S2).

### 2.8 Frequency characteristics of semi-simulated timeseries

Fourier transforms were performed on the original overlap timeseries and each of the eight semi-simulated overlap timeseries to compare the resulting frequency characteristics. Fourier transforms were performed separately for each participant and each run using only the 3T runs (8 per individual) for ease of comparison due to matched TR and timeseries length. Resulting power spectra were normalized to a maximum of 1 for ease of comparison.

### 2.9 Hidden Markov Modeling

Hidden Markov Modeling (HMM) was used to decompose the input timeseries (comprising the three regions or ‘channels’ of N1, N2, and O timeseries) into a finite number of states, where each state is a multivariate Gaussian distribution modeled by the mean (Vidaurre et al., 2017). HMM offers a valuable source of comparison of semi-simulated timeseries in addition to simple timeseries correlations because it can detect more complex dynamic states of network interaction and because it is insensitive to phase shifts which may arise from mismatches in network order for the switching hypothesis timeseries. The state covariance was modeled as one full covariance matrix for all states. The HMM was inferred separately for each participant, treating the 12 runs as separate ‘segments’ within each participant. In each participant, the HMM was repeated to test multiple options for the maximum number of states (2, 3, 4), which were chosen to range from one fewer to one more than the number of input regions (3 ‘channels’ including N1, N2, and O timeseries). As HMM is non-deterministic, we tested the stability of the HMM inference for each participant and for each maximum number of states by repeating the HMM five times and calculating the gamma similarity between each pair of repeats. The gamma similarity measures overlap between the state probabilities after optimally reordering the states (Quinn et al., 2018).

The above HMM estimation procedure was performed separately for each of the 20 network pairs using as input the original timeseries N1 and N2 in all cases, along with the original timeseries for O and separately for each of the eight semi-simulated versions of O (switch 50 TRs, switch 25 TRs, switch 10 TRs, max switching, linear additive coupling, nonlinear multiplicative coupling, spatial random mixture, spatial interdigitation). To compare the HMM state means between the original and semi-simulated versions, the intraclass correlation coefficient of absolute agreement (McGraw & Wong, 1996) was calculated after optimally reordering the states.

## 3. Results

### 3.1 PROFUMO mode maps

Out of the total of 20 modes, 12 met the test-retest and subject-group criteria to be considered for further analyses (see section 2.3). The 12 modes (Fig. 3) covered well-known occipital (visual), pericentral (somatomotor), dorsal frontoparietal (attention), lateral frontoparietal (control), and medial frontoparietal (default) networks (Uddin et al., 2019). These 12 modes were highly replicable across different dimensionalities of PROFUMO (Supplementary Fig. S3). Individual participants had on average 1.4 missing modes (range 0-4; Supplementary Fig. S4). Group maps were stable across the group-PROFUMO using all data and group-PROFUMO using only 12 runs to match subject analyses, albeit with lower weights in the 12-run results reflecting the reduction in SNR (Fig 4 top row). Mode maps for two example participants derived from single-subject PROFUMO reveal detailed individual specific organization that closely matches maps from the same participants derived from the classic PROFUMO group analysis (Fig. 4 middle and bottom rows). The example participants were chosen as non-twin individuals with a complete set of 12 modes and are representative in terms of all other indices (as shown by highlighting the blue and red data points in Figures 4, 5, 7, S4, S6, S7, S8). Subject-specific networks and twin comparisons are shown in Supplementary Fig S5.

**Figure 3:**
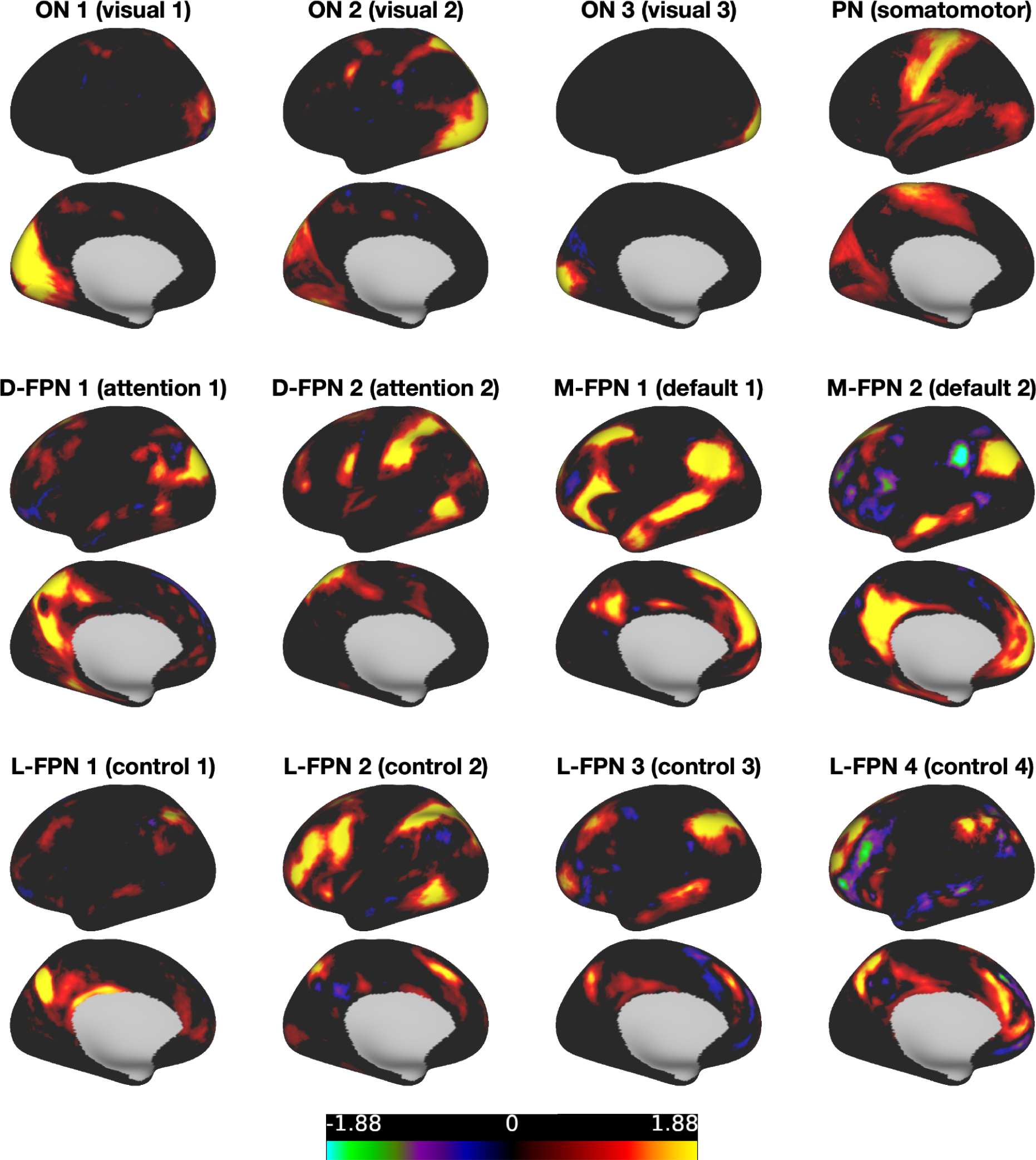
PROFUMO mode (i.e., “network”) maps derived from group PROFUMO using all 12 runs for all 20 participants. For naming purposes, modes were spatially mapped onto the Yeo-7 parcellation (Yeo et al., 2011) and we followed the naming convention suggested in (Uddin et al., 2019). FPN = frontoparietal network; L = lateral; D = dorsal; M = medial; ON = occipital network; PN = pericentral network. Figures illustrate networks on the left hemisphere. All 12 networks showed strong bilateral symmetry.

**Figure 4:**
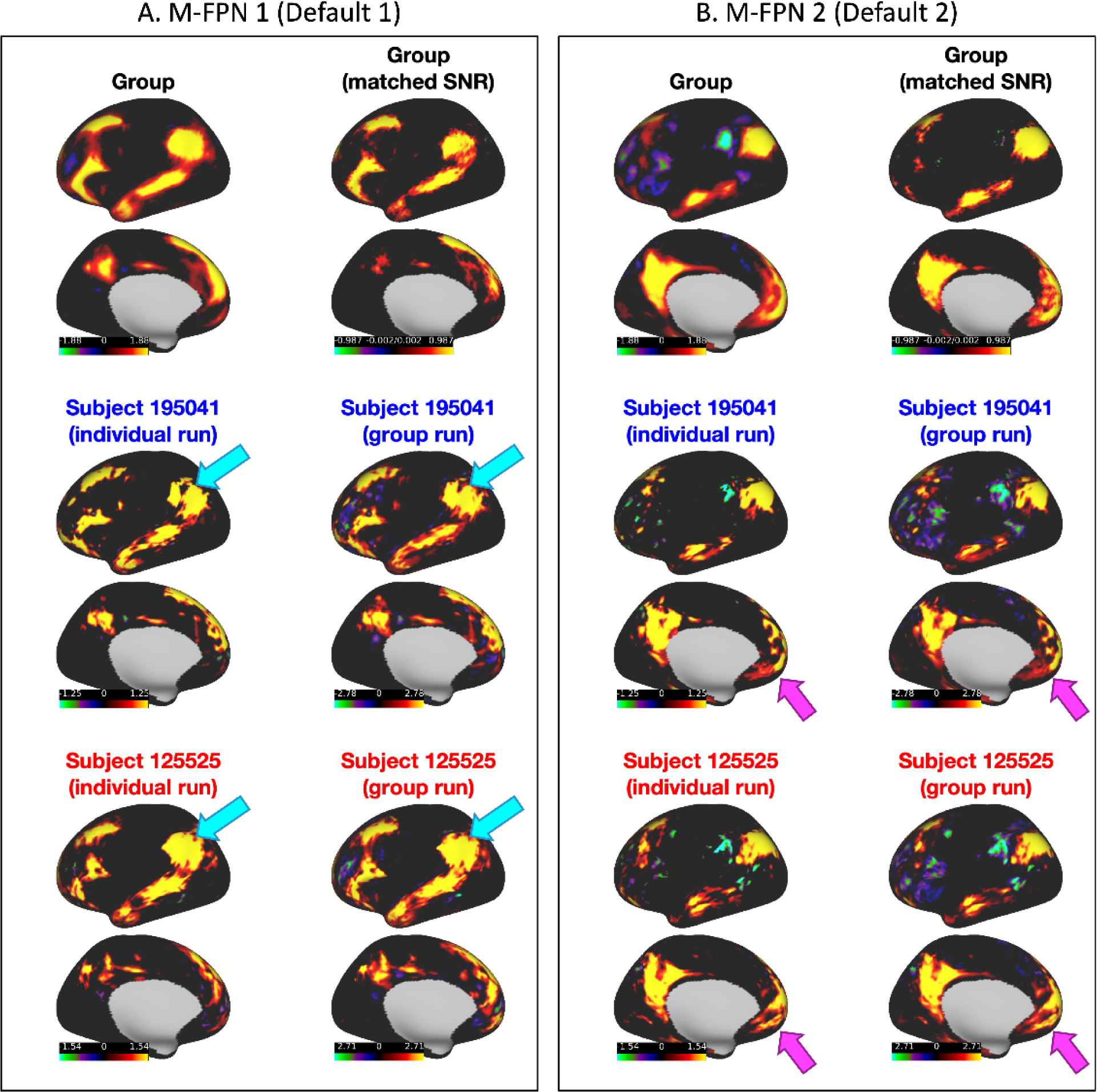
Comparison of two modes across different PROFUMO runs. A) M-FPN 1 (default 1). B) M-FPN 2 (default 2). Results display the matched mode from: classic group PROFUMO, classic group PROFUMO using only 12 scans PROFUMO runs, single-subject PROFUMO for two separate example participants, and subject-specific estimates derived from classic group PROFUMO (same two example participants). Results reveal individual differences between single-subject PROFUMO results that accurately match the estimates derived from classic group PROFUMO, confirming that PROFUMO can be used to estimate network organization using only data from a single subject. For example, the cyan arrows in (A) point to a ‘hole’ in the M-FPN 1 network that is consistently observed in subject 195041 and not in subject 125525, and the magenta arrows in (B) point to reproducible but subject-specific frontal patterns in the M-FPN 2 network.

### 3.2 PROFUMO mode stability

An initial objective was to determine whether PROFUMO can reliably estimate resting state maps using only data from a single participant. Fig. 5 shows that the highest similarity occurs for single-subject PROFUMO spatial maps (test-retest reliability and within-subject similarity in columns 1&2, respectively; mean r=0.80±0.18); the next highest similarity occurs for twin correlations (Fig 5 columns 3&4; mean r=0.70±0.14); and the lowest similarity occurs between non-twin participants (Fig 5 columns 5&6; mean r=0.59±0.15). Each similarity pattern shows a ‘tail’ of networks having lower stability (Fig. 5), which on further inspection appeared to be distributed across participants and across networks. When all modes are included (i.e., not only the 12 selected modes and not removing ‘missing’ individual modes), these tails are further expanded but the findings described above regarding comparisons between columns remain evident (see supplementary figure S6). Single-subject PROFUMO spatial maps derived using the 12 runs for the participant (i.e., not including data from other participants) were also highly similar to the estimated subject maps derived from the group data informed by all 20 subjects with 12 runs each (Fig 5 column 2; mean group-subject r=0.86±0.11), suggesting reasonable correspondence at least at this dimensionality. Spatial maps estimates informed by group data achieved slightly higher similarity compared to maps estimated from individual runs (Fig 5 column 4 > 3 and column 6 > 5), revealing the impact of the group prior when performing classic hierarchical PROFUMO. Stability estimates for second order statistics, including temporal connectivity matrices and spatial overlap matrices, are shown in Supplementary Figures S7 and S8.

**Figure 5:**
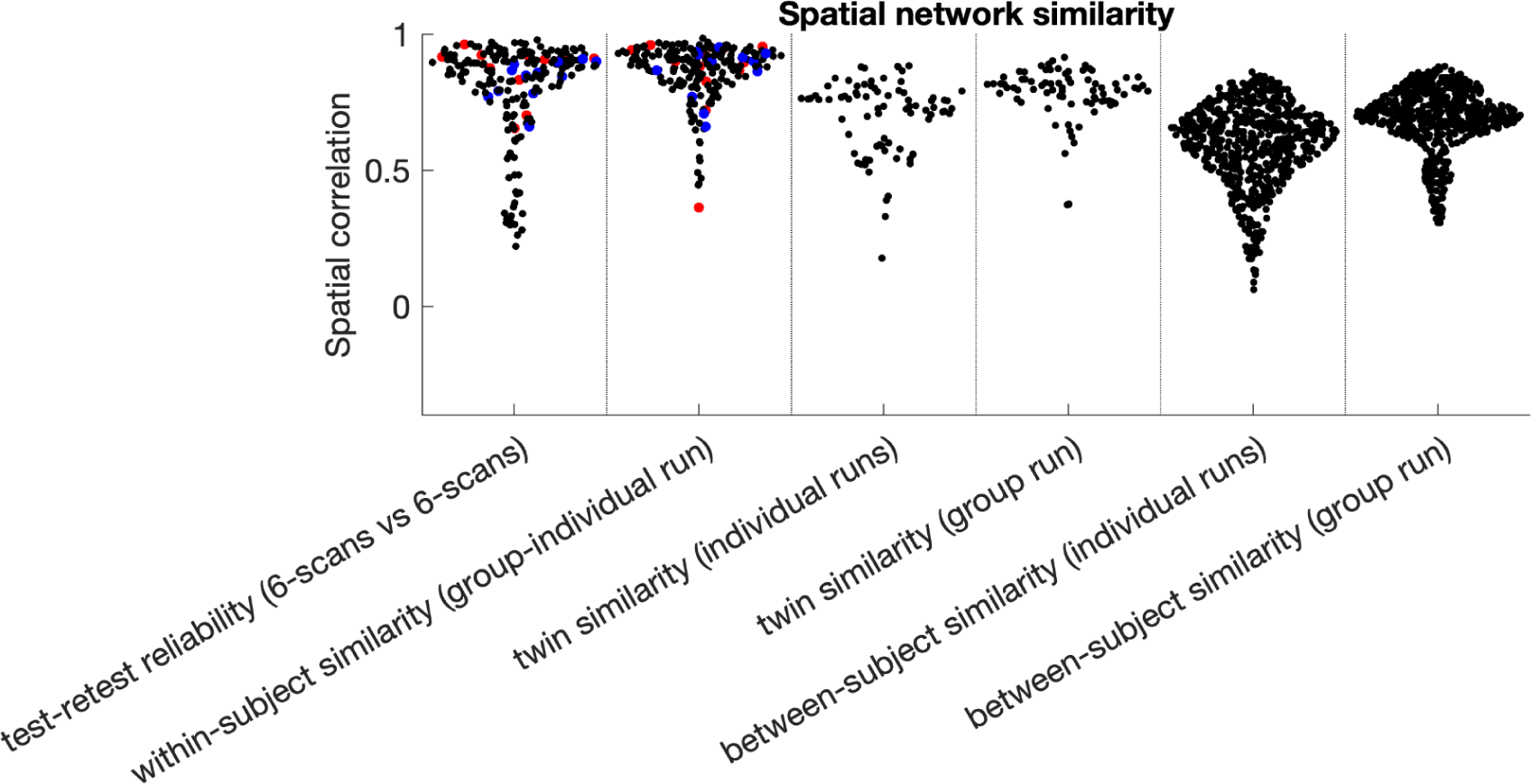
Stability of PROFUMO spatial networks. Results show high within-subject test-retest stability and similarity between individual and group estimates. Similarity within twins is lower than within individuals, but higher than between non-twin participants. Red and blue dots indicate results from the example participants used throughout this paper (separate dots are separate modes).

### 3.3 Amount of data needed to estimate single-subject modes using PROFUMO

To test how much data from an individual participant were needed to obtain good estimates from single-subject PROFUMO, we systematically varied the number of timepoints (in increments of 1/12th of the run timepoints), and compared the resulting spatial maps to the results that included all timepoints. Mean similarity across the selected (non-missing) mode maps increased when the number of timepoints were increased (Fig. 6) and was near asymptotic above ∼5000 TRs (approximately 1 hour of data per participant).

**Figure 6:**
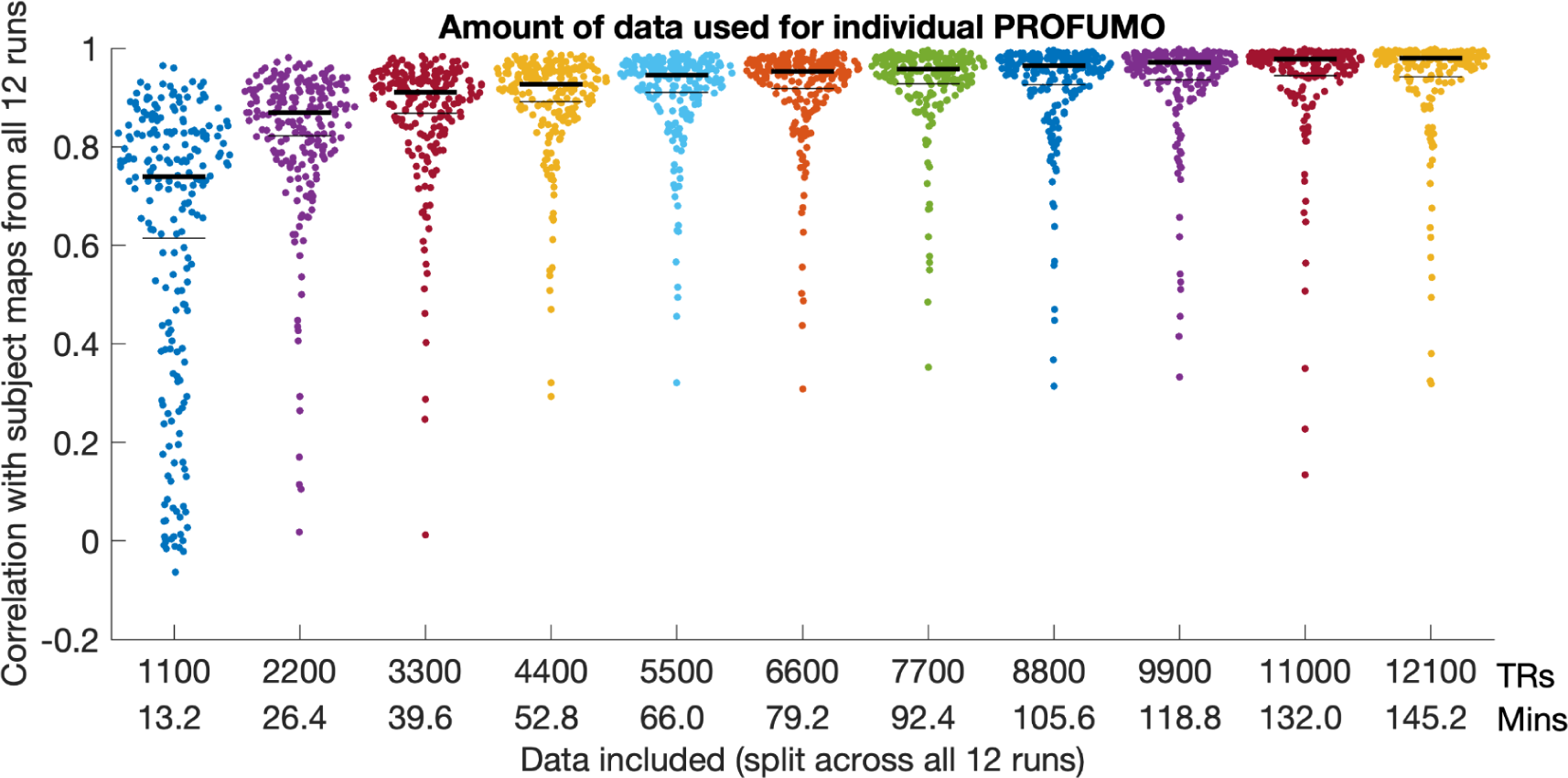
Comparison between subject maps using cut-down versions of the data relative to the subject maps obtained using the full dataset. Each dot represents a single mode-map for a single participant. Thin black lines indicate the mean and thick black lines indicate the median.

### 3.4 Spatial overlap

Consistent with previous work (Bijsterbosch et al., 2019), our results indicate substantial spatial correlation across PROFUMO modes (Fig. 7A). Figure 7A shows the spatial overlap matrix calculated by correlating pairs of weighted PROFUMO group spatial maps. Notably, spatial overlap is ignored in the find-the-biggest overview in Supplementary Fig. S5, in which vertices are assigned to a single PROFUMO mode with the highest spatial weight to enable concise visualization of all subject maps. Spatial overlap was primarily localized in the lateral parietal cortex and posterior cingulate - precuneus regions (Fig. 7C&D), consistent with prior work (Bijsterbosch et al., 2019). Spatial overlap maps for two example participants are shown in Fig. 7C and D; Supplementary Figure S9 shows maps for all participants and across twin pairs, and Supplementary Figure S1 shows maps for the selected set of 20 2-network overlap pairs.

**Figure 7:**
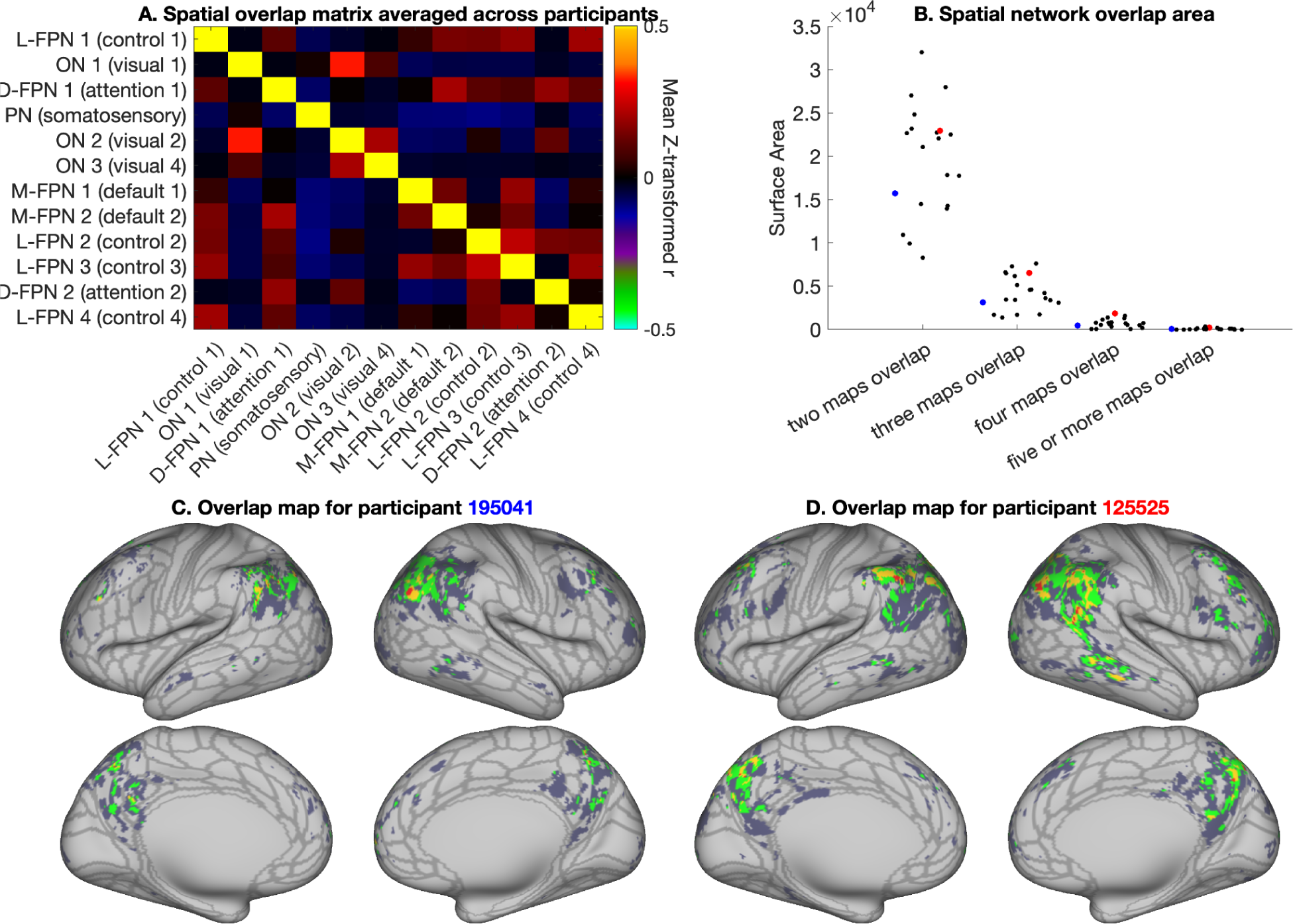
Overview of spatial overlap. A) Group average spatial overlap matrix, showing pairwise correlations between spatial maps. B) Number of vertices with 2, 3, 4, 5+ overlapping networks for each individual. Red and blue dots indicate results from the examples participants used throughout this paper. C and D) Overlap maps for the same two example participants shown in Fig. 4 (overlap maps for all participants can be found in supplementary Figure S9). For reference, the borders of the HCP_MMP1.0 cortical parcellation from (Glasser et al., 2016) are shown in gray in figures C and D.

### 3.5 Semi-simulated data for 2-network overlap

We derived semi-simulated versions of the overlap timeseries based on combinations of network 1 and network 2 timeseries (Fig. 2C,D,E). We assessed how similar the semi-simulated timeseries for the overlap region were, compared to the original overlap timeseries (O) by estimating the Pearson’s correlation coefficient separately for each network pair, each subject and each run. The highest similarity was observed for the linear additive coupling hypothesis, which achieved a median correlation of 0.783 (Fig. 8). This result was significantly different from the next highest correlation (median of 0.767), observed for the nonlinear multiplicative coupling hypothesis (T=6.3, p=3.2*10^−10^, estimated after z-transformation of the correlation values).

**Figure 8:**
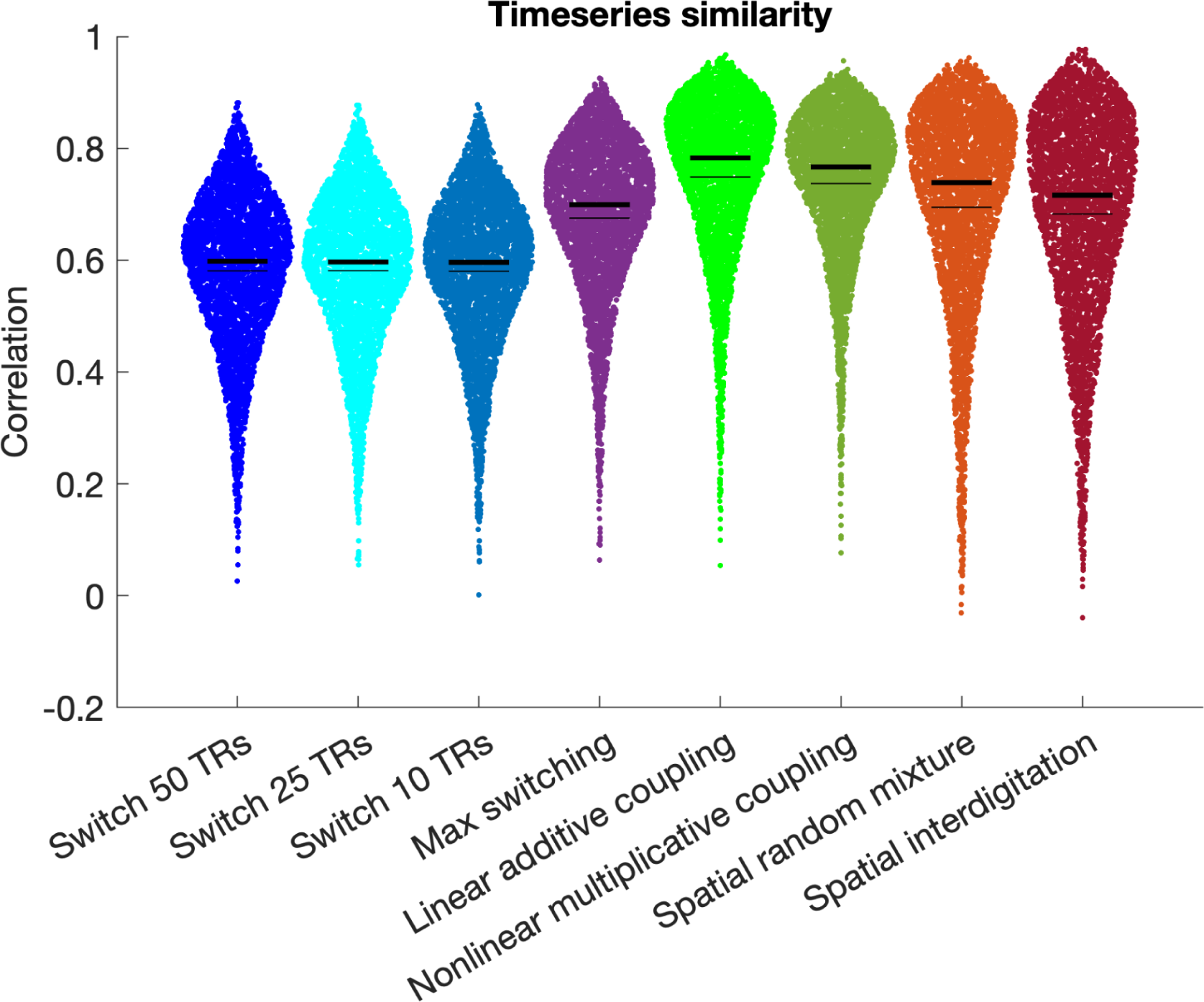
Correlations between the true overlap timeseries and different versions of the semi-simulated overlap timeseries. Thin black lines indicate the mean and thick black lines indicate the median. Highest similarity was observed for the linear additive coupling semi-simulated overlap timeseries in bright green. Data were combined across all network pairs and all participants, see supplementary figure S10 for separate figures per network pair.

Across the 20 different network pairs, the linear additive semi-simulated timeseries achieved the highest correlation for 16 network pairs, and the nonlinear multiplicative semi-simulated timeseries achieved the highest correlation for the remaining 4 network pairs (see Supplementary Fig. S10). Notably, each of the 4 network pairs with highest correlations for the nonlinear multiplicative semi-simulated timeseries involved the L-FPN 1 (paired with M-FPN 2, L-FPN 2, L-FPN 3, and L-FPN 4 respectively). The L-FPN 1 network includes a specific subregion (‘POS2’) of the parietal-occipital sulcus that is highly distinctive from neighboring regions in its myelin, thickness, connectivity, and task activation (Glasser et al., 2016). As such, the L-FPN 1 network (and POS2 area in particular) warrant future research into their distinctive features, including nonlinear origins of network overlap.

Frequency characteristics were highly similar between the original overlap timeseries and all semi-simulated timeseries (see supplementary figure S11), although spatial interdigitation timeseries exhibited a slightly raised tail and the temporal switching hypothesis (every 10 TRs) has somewhat increased power at 0.1 Hz.

### 3.6 Hidden Markov Modeling

Out of the semi-simulated results, spatial random mixture achieved the highest absolute agreement of HMM state means (mean 0.79, median 0.92), followed by spatial interdigitation (mean 0.76, median 0.88), and linear additive coupling (mean 0.75, median 0.85). HMM stability and mean similarity were consistently higher for the 2-state solution compared to the 3 and 4-state solutions (supplementary figure S12). Results separated per network pair are in Supplementary Figures S13, S14, and S15. Taken together, the HMM results provide support for the spatial mixing hypothesis despite overall lower timeseries correlations observed in Fig. 8, whilst also supporting high state similarity for linear additive coupling.

## Discussion

Our first aim was to determine whether weighted resting state spatial networks can be robustly derived from single-subject data without sacrificing correspondence. The results revealed twelve resting state networks with high test-retest reliability and remarkably good incidental (i.e., non-modeled) correspondence with group-informed network estimates (Fig. 4). Notably, the test-retest reliability of these individual-specific network maps (cosine similarity ∼0.8 based on Fig. 10A in (Harrison et al., 2020)) matches or exceeds many previous estimates of reliability of resting state metrics (e.g., average intraclass correlation 0.29 for temporal functional connectivity from meta analysis across 25 studies; (Noble et al., 2019)) (Andellini et al., 2015; Braun et al., 2012; Dutt et al., 2021; Fiecas et al., 2013; Liao et al., 2013; Nemani & Lowe, 2021; Noble et al., 2019; Termenon et al., 2016; J. Wang et al., 2017; J.-H. Wang et al., 2011; Yang et al., 2021). These findings support the possibility of future personalized psychiatry approaches where data from an individual would be separately analyzed and compared to a reference cohort to inform clinical decision making. It is possible that increasing the dimensionality to extract more finer-grained resting state networks (beyond the 12 selected networks out of 20 used here) may reduce test-retest reliability and correspondence. However, we showed high stability of the selected networks across higher dimensionalities of PROFUMO (Supplementary Fig. S1), and previous work in networks defined with independent component analysis reported good reliability up to at least 150 networks (Ma & MacDonald, 2021). Importantly, the functional organization of the brain can meaningfully be studied at multiple levels of complexity along its organizational hierarchy (Bijsterbosch et al., 2021). Here, we specifically chose to investigate the organization of macroscale networks (i.e., low dimensionality) because it describes functional organization in relation to widely studied canonical networks that are consistently observed across datasets, analysis methods, and states (Smith et al., 2009; Uddin et al., 2019; Yeo et al., 2011).

Our second aim was to determine the degree of spatially overlapping network organization. Our findings confirm the presence of extensive and stable network overlap in networks estimated from single-subject data. Patterns of spatially overlapping network organization vary extensively across individuals (Fig. 7 & Supplementary Fig. S6), and previous work has shown that individual differences in these overlap patterns are strongly associated with behavior (Bijsterbosch et al., 2019). As such, the lower dimensional spatial overlap matrix (Fig. 7A) may provide a key summary measure of behaviorally-relevant aspects of spatial organization, while reducing the multiple-comparison burden of vertex-wise analysis of spatial organization.

Spatial overlap between individual-specific weighted resting state network maps also offers a complementary approach to investigate hub regions. As opposed to traditional hub identification methods that rely on temporal correlations, our weighted network approach emphasizes the role of shared brain regions as part of the spatial organization of functional networks. It is currently unclear whether these distinct temporal versus spatial definitions of hub regions have dissociable or shared neurobiological implications. Some evidence suggests that spatial networks and hubs may maintain functional integrity over time, potentially for homeostatic purposes (Laumann & Snyder, 2021), whereas flexible hubs that can rapidly change their temporal connectivity may provide coordination and switching functions to support cognitive control (Cocuzza et al., 2020; Cole et al., 2013; Gordon et al., 2018). Consistent with this hypothesis, we have shown that spatial overlap was more strongly associated with stable trait-like behavior (Bijsterbosch et al., 2018), whereas temporal correlations tracked transitions between sensorimotor states (Harrison et al., 2020). However, our prior work also suggests that effects of spatial network organization and overlap are observed as temporal correlation estimates when unaccounted for in the parcellation (Bijsterbosch et al., 2018, 2019), which may indicate that the distinction between spatial and temporal hubs could be purely analytical, driven by differences in model cost functions and priors, without distinct neurobiological interpretations. Hence, spatial network overlap and temporally strongly connected nodes may serve distinct functional purposes (e.g., homeostasis versus switching), or may represent alternative analytical estimates of the same underlying neural phenomenon. Further work is needed to gain insight into spatial versus temporal hub definitions and their neurobiological functions.

Our third aim was to systematically test different hypotheses (spatial mixture, dynamic switching, and coupling hypotheses; Fig. 1) regarding the nature of brain regions in which multiple resting state networks appear to overlap. Our findings supported the linear additive coupling hypothesis (Figs. 8&9). Given that regions of network overlap appear to be actively engaged in processing information from all contributing networks, this suggests that network overlap may play an important role in the integration of information across multiple brain systems. Alternatively, the linear additive hypothesis may suggest that multiple networks may coexist within overlap regions without influencing one another. Specifically, the combined signal may indicate that both networks behave as they do in non-overlap regions without any cross-network modulation or integration. The nonlinear hypothesis, on the other hand, requires cross-network integration. Prior work has reported that nonlinear binding between multiple task conditions in conjunction hubs (i.e., brain regions that selectively integrate activations) was essential for predicting task activation patterns from functional connectivity data (Ito et al., 2022). Intriguingly, the timeseries correlation results (Supplementary Figure S10) offered some support for the nonlinear multiplicative hypothesis for network pairs involving the L-FPN 2, which may indicate the presence of network interaction. Importantly, it remains a possibility that spatial mixing and/or dynamic switching may occur at finer spatial and temporal resolutions that cannot be resolved using resting state MRI data. Furthermore, it is possible that temporally lagged correlation structure may play a role in regions of spatial overlap, which can be challenging to accurately discern using functional MRI data (Smith et al., 2011) but may be feasible in some situations using deconvolution approaches (Mill et al., 2017). Although beyond the scope of the current paper, we plan to test spatial mixing, dynamic switching, and temporally lagged hypotheses at sub-MRI scales in future research using invasive recording techniques in non-human primates.

**Figure 9:**
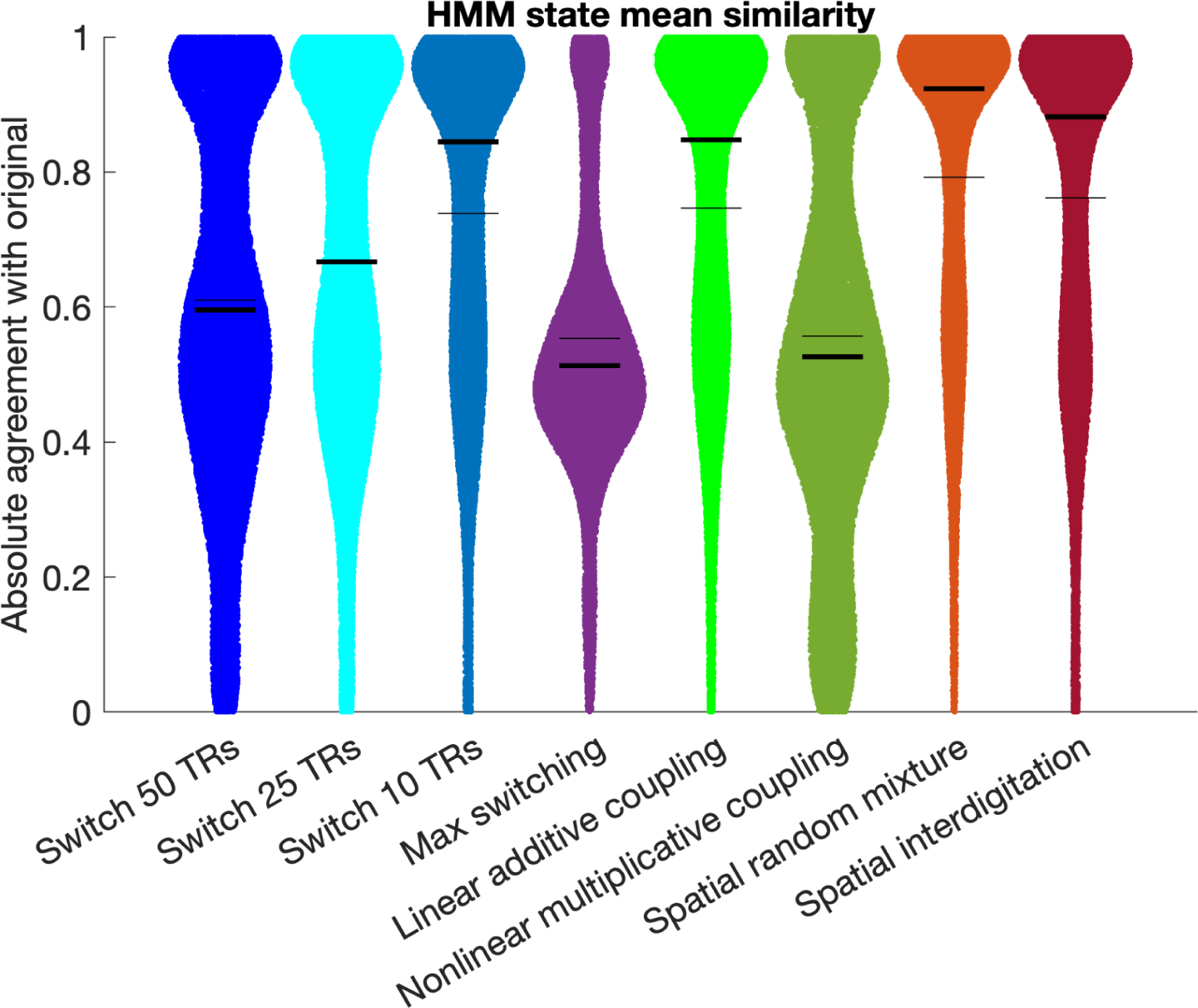
Absolute agreement between the HMM state means estimated when using the true overlap timeseries compared to different versions of the semi-simulated overlap timeseries. Thin black lines indicate the mean and thick black lines indicate the median. Data were combined across all network pairs, all participants, and all state solutions. Additional results separated for 2, 3, and 4 state solutions are in Supplementary Figure S12 and separate results per network pair are shown in Supplementary Figures S13 (2-state), S14 (3-state), and S15 (4-state).

Despite offering novel insights into weighted resting state networks estimated within individual participants, this work also has several limitations. First, we investigated network organization at a relatively coarse level by setting the PROFUMO dimensionality to 20 and investigating 12 resulting networks of high stability. Notably, the 12 networks under investigations were obtained with very high stability across a range of PROFUMO dimensionalities (Supplementary Fig. S1). Nevertheless, network overlap may behave differently at higher dimensions of network decomposition (Farahibozorg et al., 2021). Second, our vertex assignments to network 1, network 2, and overlap regions involved thresholding, which is necessarily a simplification of the weighted network organization. However, the strict threshold for exclusion and inclusion of vertices enforced a relatively conservative definition of overlap that was necessary to make the results interpretable. Importantly, we recommend the use of the unthresholded spatial overlap matrix (Fig. 7A) for brain-behavior investigations of network overlap. Third, the estimation of network overlap, whether based on weighted or thresholded maps, interacts with the contrast-to-noise ratio (CNR) of the data, which varies across the cortex and is higher in the lateral parietal region. As such, potential future efforts to develop a conclusive map of network overlap in the brain should perform careful scaling/normalization to remove any bias resulting from CNR variability across the cortex. Furthermore, individual difference studies should match the amount of data used across individuals to avoid biased estimates of network overlap that may arise from CNR variability across individuals. Fourth, our findings indicating support for the linear additive coupling hypothesis are consistent with the underlying outer product model used in PROFUMO (and similar methods such as ICA). In future work, it will be of interest to test mechanistic hypotheses of network overlap using alternative approaches to define the overlap region that do not rely on the outer product model. Notably, alternative tools for the definition of network overlap (such as (Karahanoğlu & Van De Ville, 2015; Li et al., 2017)) might render different overlap results and might have different reliability and data-needs.

Taken together, we showed that weighted resting state networks derived from single-subject data are stable, correspond closely to group-informed networks, and capture overlapping network organization, and are therefore important targets for clinical biomarker research. We also showed that overlapping network organization is indicative of linear coupling between networks, providing a mechanistic hypothesis for the functional role of these regions.

## Ethics

Secondary data analysis of HCP data for this project was covered under IRB 202105086.

## Data and Code Availability

The data used in this paper are freely available from https://db.humanconnectome.org/ (requires free registration). Hidden Markov Modeling was implemented using the HMM-MAR Matlab toolbox: https://github.com/OHBA-analysis/HMM-MAR/wiki. All code for this paper is available on GitHub: https://github.com/JanineBijsterbosch/Individual_PROFUMO.

## Author Contributions

JDB conceived of the study, performed all analyses, and wrote the manuscript. SRF provided support regarding the PROFUMO algorithm. MFG and DVE helped with HCP aspects and interpretation. LHS contributed to the hypothesis generation and funding. MWW and SMS helped with the analysis design and interpretation. All authors reviewed the manuscript.

## References

1. Andellini, M., Cannatà, V., Gazzellini, S., Bernardi, B., & Napolitano, A. (2015). Test-retest reliability of graph metrics of resting state MRI functional brain networks: A review. Journal of Neuroscience Methods, 253, 183–192. 10.1016/j.jneumeth.2015.05.020

2. Bertolero, M. A., Yeo, B. T. T., Bassett, D. S., & D’Esposito, M. (2018). A mechanistic model of connector hubs, modularity and cognition. Nature Human Behaviour, 2(10), 765–777. 10.1038/s41562-018-0420-6

3. Bijsterbosch, J. D., Beckmann, C. F., Woolrich, M. W., Smith, S. M., & Harrison, S. J. (2019). The relationship between spatial configuration and functional connectivity of brain regions revisited. eLife, 8. 10.7554/eLife.44890

4. Bijsterbosch, J. D., Harrison, S. J., Jbabdi, S., Woolrich, M., Beckmann, C., Smith, S., & Duff, E. P. (2020). Challenges and future directions for representations of functional brain organization. Nature Neuroscience, 1–12. 10.1038/s41593-020-00726-z

5. Bijsterbosch, J. D., Valk, S. L., Wang, D., & Glasser, M. F. (2021). Recent developments in representations of the connectome. NeuroImage, 243, 118533. 10.1016/j.neuroimage.2021.118533

6. Bijsterbosch, J. D., Woolrich, M. W., Glasser, M. F., Robinson, E. C., Beckmann, C. F., Van Essen, D. C., Harrison, S. J., & Smith, S. M. (2018). The relationship between spatial configuration and functional connectivity of brain regions. eLife, 7. 10.7554/eLife.32992

7. Blazquez Freches, G., Haak, K. V., Bryant, K. L., Schurz, M., Beckmann, C. F., & Mars, R. B. (2020). Principles of temporal association cortex organisation as revealed by connectivity gradients. Brain Structure & Function, 225(4), 1245–1260. 10.1007/s00429-020-02047-0

8. Braga, R. M., & Buckner, R. L. (2017). Parallel Interdigitated Distributed Networks within the Individual Estimated by Intrinsic Functional Connectivity. Neuron, 95(2), 457–471.e5. 10.1016/j.neuron.2017.06.038

9. Braga, R. M., Van Dijk, K. R. A., Polimeni, J. R., Eldaief, M. C., & Buckner, R. L. (2019). Parallel distributed networks resolved at high resolution reveal close juxtaposition of distinct regions. Journal of Neurophysiology, 121(4), 1513–1534. 10.1152/jn.00808.2018

10. Braun, U., Plichta, M. M., Esslinger, C., Sauer, C., Haddad, L., Grimm, O., Mier, D., Mohnke, S., Heinz, A., Erk, S., Walter, H., Seiferth, N., Kirsch, P., & Meyer-Lindenberg, A. (2012). Test–retest reliability of resting-state connectivity network characteristics using fMRI and graph theoretical measures. NeuroImage, 59(2), 1404–1412. 10.1016/j.neuroimage.2011.08.044

11. Buckner, R. L., Sepulcre, J., Talukdar, T., Krienen, F. M., Liu, H., Hedden, T., Andrews-Hanna, J. R., Sperling, R. A., & Johnson, K. A. (2009). Cortical hubs revealed by intrinsic functional connectivity: mapping, assessment of stability, and relation to Alzheimer’s disease. The Journal of Neuroscience: The Official Journal of the Society for Neuroscience, 29(6), 1860–1873. 10.1523/JNEUROSCI.5062-08.2009

12. Cocuzza, C. V., Ito, T., Schultz, D., Bassett, D. S., & Cole, M. W. (2020). Flexible Coordinator and Switcher Hubs for Adaptive Task Control. The Journal of Neuroscience: The Official Journal of the Society for Neuroscience, 40(36), 6949–6968. 10.1523/JNEUROSCI.2559-19.2020

13. Cole, M. W., Reynolds, J. R., Power, J. D., Repovs, G., Anticevic, A., & Braver, T. S. (2013). Multi-task connectivity reveals flexible hubs for adaptive task control. Nature Neuroscience, 16(9), 1348–1355. 10.1038/nn.3470

14. Dutt, R. K., Hannon, K., Easley, T. O., Griffis, J. C., Zhang, W., & Bijsterbosch, J. D. (2021). Mental health in the UK Biobank: A roadmap to self-report measures and neuroimaging correlates. Human Brain Mapping. 10.1002/hbm.25690

15. Farahibozorg, S.-R., Bijsterbosch, J. D., Gong, W., Jbabdi, S., Smith, S. M., Harrison, S. J., & Woolrich, M. W. (2021). Hierarchical modelling of functional brain networks in population and individuals from big fMRI data. NeuroImage, 118513. 10.1016/j.neuroimage.2021.118513

16. Fiecas, M., Ombao, H., van Lunen, D., Baumgartner, R., Coimbra, A., & Feng, D. (2013). Quantifying temporal correlations: A test–retest evaluation of functional connectivity in resting-state fMRI. NeuroImage, 65, 231–241. 10.1016/j.neuroimage.2012.09.052

17. Glasser, M. F., Coalson, T. S., Robinson, E. C., Hacker, C. D., Harwell, J., Yacoub, E., Ugurbil, K., Andersson, J., Beckmann, C. F., Jenkinson, M., Smith, S. M., & Van Essen, D. C. (2016). A multi-modal parcellation of human cerebral cortex. Nature, 536(7615), 171–178. 10.1038/nature18933

18. Glasser, M. F., Sotiropoulos, S. N., Wilson, J. A., Coalson, T. S., Fischl, B., Andersson, J. L., Xu, J., Jbabdi, S., Webster, M., Polimeni, J. R., Van Essen, D. C., Jenkinson, M., & WU-Minn HCP Consortium. (2013). The minimal preprocessing pipelines for the Human Connectome Project. NeuroImage, 80, 105–124. 10.1016/j.neuroimage.2013.04.127

19. Gordon, E. M., Laumann, T. O., Adeyemo, B., Gilmore, A. W., Nelson, S. M., Dosenbach, N. U. F., & Petersen, S. E. (2017). Individual-specific features of brain systems identified with resting state functional correlations. NeuroImage, 146, 918–939. 10.1016/j.neuroimage.2016.08.032

20. Gordon, E. M., Laumann, T. O., Adeyemo, B., & Petersen, S. E. (2017). Individual Variability of the System-Level Organization of the Human Brain. Cerebral Cortex, 27(1), 386–399. 10.1093/cercor/bhv239

21. Gordon, E. M., Laumann, T. O., Gilmore, A. W., Newbold, D. J., Greene, D. J., Berg, J. J., Ortega, M., Hoyt-Drazen, C., Gratton, C., Sun, H., Hampton, J. M., Coalson, R. S., Nguyen, A. L., McDermott, K. B., Shimony, J. S., Snyder, A. Z., Schlaggar, B. L., Petersen, S. E., Nelson, S. M., & Dosenbach, N. U. F. (2017). Precision Functional Mapping of Individual Human Brains. Neuron, 95(4), 791–807.e7. 10.1016/j.neuron.2017.07.011

22. Gordon, E. M., Lynch, C. J., Gratton, C., Laumann, T. O., Gilmore, A. W., Greene, D. J., Ortega, M., Nguyen, A. L., Schlaggar, B. L., Petersen, S. E., Dosenbach, N. U. F., & Nelson, S. M. (2018). Three Distinct Sets of Connector Hubs Integrate Human Brain Function. Cell Reports, 24(7), 1687–1695.e4. 10.1016/j.celrep.2018.07.050

23. Gratton, C., Kraus, B. T., Greene, D. J., Gordon, E. M., Laumann, T. O., Nelson, S. M., Dosenbach, N. U. F., & Petersen, S. E. (2020). Defining Individual-Specific Functional Neuroanatomy for Precision Psychiatry. Biological Psychiatry, 88(1), 28–39. 10.1016/j.biopsych.2019.10.026

24. Gratton, C., Laumann, T. O., Nielsen, A. N., Greene, D. J., Gordon, E. M., Gilmore, A. W., Nelson, S. M., Coalson, R. S., Snyder, A. Z., Schlaggar, B. L., Dosenbach, N. U. F., & Petersen, S. E. (2018). Functional Brain Networks Are Dominated by Stable Group and Individual Factors, Not Cognitive or Daily Variation. Neuron, 98(2), 439–452.e5. 10.1016/j.neuron.2018.03.035

25. Griffanti, L., Salimi-Khorshidi, G., Beckmann, C. F., Auerbach, E. J., Douaud, G., Sexton, C. E., Zsoldos, E., Ebmeier, K. P., Filippini, N., Mackay, C. E., Moeller, S., Xu, J., Yacoub, E., Baselli, G., Ugurbil, K., Miller, K. L., & Smith, S. M. (2014). ICA-based artefact removal and accelerated fMRI acquisition for improved resting state network imaging. NeuroImage, 95, 232–247. 10.1016/j.neuroimage.2014.03.034

26. Haak, K. V., Marquand, A. F., & Beckmann, C. F. (2018). Connectopic mapping with resting-state fMRI. NeuroImage, 170, 83–94. 10.1016/j.neuroimage.2017.06.075

27. Harrison, S. J., Bijsterbosch, J. D., Segerdahl, A. R., Fitzgibbon, S. P., Farahibozorg, S.-R., Duff, E. P., Smith, S. M., & Woolrich, M. W. (2020). Modelling subject variability in the spatial and temporal characteristics of functional modes. NeuroImage, 222, 117226. 10.1016/j.neuroimage.2020.117226

28. Harrison, S. J., Woolrich, M. W., Robinson, E. C., Glasser, M. F., Beckmann, C. F., Jenkinson, M., & Smith, S. M. (2015). Large-scale probabilistic functional modes from resting state fMRI. NeuroImage, 109, 217–231. 10.1016/j.neuroimage.2015.01.013

29. Hutchison, R. M., Womelsdorf, T., Allen, E. A., Bandettini, P. A., Calhoun, V. D., Corbetta, M., Della Penna, S., Duyn, J. H., Glover, G. H., Gonzalez-Castillo, J., Handwerker, D. A., Keilholz, S., Kiviniemi, V., Leopold, D. A., de Pasquale, F., Sporns, O., Walter, M., & Chang, C. (2013). Dynamic functional connectivity: promise, issues, and interpretations. NeuroImage, 80, 360–378. 10.1016/j.neuroimage.2013.05.079

30. Insel, T. R. (2014). The NIMH Research Domain Criteria (RDoC) Project: precision medicine for psychiatry. The American Journal of Psychiatry, 171(4), 395–397. 10.1176/appi.ajp.2014.14020138

31. Ito, T., Yang, G. R., Laurent, P., Schultz, D. H., & Cole, M. W. (2022). Constructing neural network models from brain data reveals representational transformations linked to adaptive behavior. Nature Communications, 13(1), 673. 10.1038/s41467-022-28323-7

32. Karahanoğlu, F. I., & Van De Ville, D. (2015). Transient brain activity disentangles fMRI resting-state dynamics in terms of spatially and temporally overlapping networks. Nature Communications, 6, 7751. 10.1038/ncomms8751

33. Kong, R., Li, J., Orban, C., Sabuncu, M. R., Liu, H., Schaefer, A., Sun, N., Zuo, X.-N., Holmes, A. J., Eickhoff, S. B., & Yeo, B. T. T. (2019). Spatial Topography of Individual-Specific Cortical Networks Predicts Human Cognition, Personality, and Emotion. Cerebral Cortex, 29(6), 2533–2551. 10.1093/cercor/bhy123

34. Kuhn, H. W. (1955). The Hungarian method for the assignment problem. Naval Research Logistics Quarterly, 2(1-2), 83–97. 10.1002/nav.3800020109

35. Laumann, T. O., Gordon, E. M., Adeyemo, B., Snyder, A. Z., Joo, S. J., Chen, M.-Y., Gilmore, A. W., McDermott, K. B., Nelson, S. M., Dosenbach, N. U. F., Schlaggar, B. L., Mumford, J. A., Poldrack, R. A., & Petersen, S. E. (2015). Functional System and Areal Organization of a Highly Sampled Individual Human Brain. Neuron, 87(3), 657–670. 10.1016/j.neuron.2015.06.037

36. Laumann, T. O., & Snyder, A. Z. (2021). Brain activity is not only for thinking. Current Opinion in Behavioral Sciences, 40, 130–136. 10.1016/j.cobeha.2021.04.002

37. Lee, K., Lina, J.-M., Gotman, J., & Grova, C. (2016). SPARK: Sparsity-based analysis of reliable k-hubness and overlapping network structure in brain functional connectivity. NeuroImage, 134, 434–449. 10.1016/j.neuroimage.2016.03.049

38. Liao, X.-H., Xia, M.-R., Xu, T., Dai, Z.-J., Cao, X.-Y., Niu, H.-J., Zuo, X.-N., Zang, Y.-F., & He, Y. (2013). Functional brain hubs and their test–retest reliability: A multiband resting-state functional MRI study. NeuroImage, 83, 969–982. 10.1016/j.neuroimage.2013.07.058

39. Li, H., Satterthwaite, T. D., & Fan, Y. (2017). Large-scale sparse functional networks from resting state fMRI. NeuroImage, 156, 1–13. 10.1016/j.neuroimage.2017.05.004

40. Lin, Y., Ma, J., Gu, Y., Yang, S., Li, L. M. W., & Dai, Z. (2018). Intrinsic overlapping modular organization of human brain functional networks revealed by a multiobjective evolutionary algorithm. NeuroImage, 181, 430–445. 10.1016/j.neuroimage.2018.07.019

41. Marek, S., Tervo-Clemmens, B., Calabro, F. J., Montez, D. F., Kay, B. P., Hatoum, A. S., Donohue, M. R., Foran, W., Miller, R. L., Hendrickson, T. J., Malone, S. M., Kandala, S., Feczko, E., Miranda-Dominguez, O., Graham, A. M., Earl, E. A., Perrone, A. J., Cordova, M., Doyle, O.,…Dosenbach, N. U. F. (2022). Reproducible brain-wide association studies require thousands of individuals. Nature, 603(7902), 654–660. 10.1038/s41586-022-04492-9

42. Ma, Y., & MacDonald, A. W., Iii. (2021). Impact of Independent Component Analysis Dimensionality on the Test-Retest Reliability of Resting-State Functional Connectivity. Brain Connectivity, 11(10), 875–886. 10.1089/brain.2020.0970

43. McGraw, K. O., & Wong, S. P. (1996). Forming inferences about some intraclass correlation coefficients. Psychological Methods, 1(1), 30–46. 10.1037/1082-989X.1.1.30

44. Mill, R. D., Bagic, A., Bostan, A., Schneider, W., & Cole, M. W. (2017). Empirical validation of directed functional connectivity. NeuroImage, 146, 275–287. 10.1016/j.neuroimage.2016.11.037

45. Munkres, J. (1957). Algorithms for the Assignment and Transportation Problems. Journal of the Society for Industrial and Applied Mathematics, 5(1), 32–38. 10.1137/0105003

46. Najafi, M., McMenamin, B. W., Simon, J. Z., & Pessoa, L. (2016). Overlapping communities reveal rich structure in large-scale brain networks during rest and task conditions. NeuroImage, 135, 92–106. 10.1016/j.neuroimage.2016.04.054

47. Nemani, A., & Lowe, M. J. (2021). Seed-based test-retest reliability of resting state functional magnetic resonance imaging at 3T and 7T. Medical Physics, 48(10), 5756–5764. 10.1002/mp.15210

48. Noble, S., Scheinost, D., & Constable, R. T. (2019). A decade of test-retest reliability of functional connectivity: A systematic review and meta-analysis. NeuroImage, 203, 116157. 10.1016/j.neuroimage.2019.116157

49. Poldrack, R. A. (2017). Precision Neuroscience: Dense Sampling of Individual Brains. Neuron, 95(4), 727–729. 10.1016/j.neuron.2017.08.002

50. Poldrack, R. A., Laumann, T. O., Koyejo, O., Gregory, B., Hover, A., Chen, M.-Y., Gorgolewski, K. J., Luci, J., Joo, S. J., Boyd, R. L., Hunicke-Smith, S., Simpson, Z. B., Caven, T., Sochat, V., Shine, J. M., Gordon, E., Snyder, A. Z., Adeyemo, B., Petersen, S. E.,…Mumford, J. A. (2015). Long-term neural and physiological phenotyping of a single human. Nature Communications, 6, 8885. 10.1038/ncomms9885

51. Quinn, A. J., Vidaurre, D., Abeysuriya, R., Becker, R., Nobre, A. C., & Woolrich, M. W. (2018). Task-Evoked Dynamic Network Analysis Through Hidden Markov Modeling. Frontiers in Neuroscience, 12, 603. 10.3389/fnins.2018.00603

52. Robinson, E. C., Jbabdi, S., Glasser, M. F., Andersson, J., Burgess, G. C., Harms, M. P., Smith, S. M., Van Essen, D. C., & Jenkinson, M. (2014). MSM: a new flexible framework for Multimodal Surface Matching. NeuroImage, 100, 414–426. 10.1016/j.neuroimage.2014.05.069

53. Salimi-Khorshidi, G., Douaud, G., Beckmann, C. F., Glasser, M. F., Griffanti, L., & Smith, S. M. (2014). Automatic denoising of functional MRI data: combining independent component analysis and hierarchical fusion of classifiers. NeuroImage, 90, 449–468. 10.1016/j.neuroimage.2013.11.046

54. Smith, S. M., Beckmann, C. F., Andersson, J., Auerbach, E. J., Bijsterbosch, J., Douaud, G., Duff, E., Feinberg, D. A., Griffanti, L., Harms, M. P., Kelly, M., Laumann, T., Miller, K. L., Moeller, S., Petersen, S., Power, J., Salimi-Khorshidi, G., Snyder, A. Z., Vu, A. T.,…WU-Minn HCP Consortium. (2013). Resting-state fMRI in the Human Connectome Project. NeuroImage, 80, 144–168. 10.1016/j.neuroimage.2013.05.039

55. Smith, S. M., Fox, P. T., Miller, K. L., Glahn, D. C., Fox, P. M., Mackay, C. E., Filippini, N., Watkins, K. E., Toro, R., Laird, A. R., & Beckmann, C. F. (2009). Correspondence of the brain’s functional architecture during activation and rest. Proceedings of the National Academy of Sciences of the United States of America, 106(31), 13040–13045. 10.1073/pnas.0905267106

56. Smith, S. M., Miller, K. L., Salimi-Khorshidi, G., Webster, M., Beckmann, C. F., Nichols, T. E., Ramsey, J. D., & Woolrich, M. W. (2011). Network modelling methods for FMRI. NeuroImage, 54(2), 875–891. 10.1016/j.neuroimage.2010.08.063

57. Termenon, M., Jaillard, A., Delon-Martin, C., & Achard, S. (2016). Reliability of graph analysis of resting state fMRI using test-retest dataset from the Human Connectome Project. NeuroImage, 142, 172–187. 10.1016/j.neuroimage.2016.05.062

58. T Vu, A., Jamison, K., Glasser, M. F., Smith, S. M., Coalson, T., Moeller, S., Auerbach, E. J., Uğurbil, K., & Yacoub, E. (2017). Tradeoffs in pushing the spatial resolution of fMRI for the 7T Human Connectome Project. NeuroImage, 154, 23–32. 10.1016/j.neuroimage.2016.11.049

59. Uddin, L. Q., Yeo, B. T. T., & Spreng, R. N. (2019). Towards a Universal Taxonomy of Macro-scale Functional Human Brain Networks. Brain Topography. 10.1007/s10548-019-00744-6

60. Van Essen, D. C., Smith, S. M., Barch, D. M., Behrens, T. E. J., Yacoub, E., Ugurbil, K., & WU-Minn HCP Consortium. (2013). The WU-Minn Human Connectome Project: an overview. NeuroImage, 80(0), 62–79. 10.1016/j.neuroimage.2013.05.041

61. Vidaurre, D., Smith, S. M., & Woolrich, M. W. (2017). Brain network dynamics are hierarchically organized in time. Proceedings of the National Academy of Sciences of the United States of America, 114(48), 12827–12832. 10.1073/pnas.1705120114

62. Wang, D., Buckner, R. L., Fox, M. D., Holt, D. J., Holmes, A. J., Stoecklein, S., Langs, G., Pan, R., Qian, T., Li, K., Baker, J. T., Stufflebeam, S. M., Wang, K., Wang, X., Hong, B., & Liu, H. (2015). Parcellating cortical functional networks in individuals. Nature Neuroscience, 18(12), 1853–1860. 10.1038/nn.4164

63. Wang, J., Han, J., Nguyen, V. T., Guo, L., & Guo, C. C. (2017). Improving the Test-Retest Reliability of Resting State fMRI by Removing the Impact of Sleep. Frontiers in Neuroscience, 11, 249. 10.3389/fnins.2017.00249

64. Wang, J.-H., Zuo, X.-N., Gohel, S., Milham, M. P., Biswal, B. B., & He, Y. (2011). Graph theoretical analysis of functional brain networks: test-retest evaluation on short- and long-term resting-state functional MRI data. PloS One, 6(7), e21976. 10.1371/journal.pone.0021976

65. Warren, D. E., Power, J. D., Bruss, J., Denburg, N. L., Waldron, E. J., Sun, H., Petersen, S. E., & Tranel, D. (2014). Network measures predict neuropsychological outcome after brain injury. Proceedings of the National Academy of Sciences of the United States of America, 111(39), 14247–14252. 10.1073/pnas.1322173111

66. Williams, L. M. (2016). Precision psychiatry: a neural circuit taxonomy for depression and anxiety. The Lancet. Psychiatry, 3(5), 472–480. 10.1016/S2215-0366(15)00579-9

67. Yang, L., Wei, J., Li, Y., Wang, B., Guo, H., Yang, Y., & Xiang, J. (2021). Test-Retest Reliability of Synchrony and Metastability in Resting State fMRI. Brain Sciences, 12(1). 10.3390/brainsci12010066

68. Yeo, B. T. T., Krienen, F. M., Sepulcre, J., Sabuncu, M. R., Lashkari, D., Hollinshead, M., Roffman, J. L., Smoller, J. W., Zöllei, L., Polimeni, J. R., Fischl, B., Liu, H., & Buckner, R. L. (2011). The organization of the human cerebral cortex estimated by intrinsic functional connectivity. Journal of Neurophysiology, 106(3), 1125–1165. 10.1152/jn.00338.2011

