## Supplementary Materials for "Evaluating functional brain organization in individuals and identifying contributions to network overlap"

### Supplementary Information

#### Table S1: PROFUMO model details at the group level

| | | | |
|---|---|---|---|
| Group Spatial Model | Group signal component | $\mu_{vm}$<br>$p(\mu_{vm} \mid \rho_{vm} = 2) = \mathcal{N}(\mu_{vm} \mid \tau_{\mu 2}, {\gamma_{\mu 2}}^2)$<br>$p(\mu_{vm} \mid \rho_{vm} = 1) = \mathcal{N}(\mu_{vm} \mid \tau_{\mu 1}, {\gamma_{\mu 1}}^2)$<br>$p(\mu_{vm} \mid \rho_{vm} = 0) = \delta(\mu_{vm})$<br>$p(\rho_{vm}) = \prod_{i \in \{0,1,2\}} \left(\lambda_{\mu i}\right)^{[\rho_{vm} = i]}$ | Hyperprior: mixture model prior over group means (inspired by spike-slab distribution to allow sparsity, but modeled as one delta and two Gaussians to allow heavier tails)<br>Hyperparameters of first Gaussian:<br>$\square_{\mu 1}$ and $\gamma^2_{\mu 1}$<br>Hyperparameters of second Gaussian: $\square_{\mu 2}$ and $\gamma^2_{\mu 2}$ |
| | Standard deviation of signal component | $\sigma_{vm}$<br>$p(\sigma_{vm}) = \Gamma\left(\sigma_{vm}^{-2} \mid a_\sigma, b_\sigma\right)$ | Hyperprior: inverse-Gamma distribution, with hyperparameters $a_\sigma$ and $b_\sigma$ |
| | Standard deviation of noise component | $\eta_m^{(s)} \zeta_v$<br>$p(\zeta_v) = \Gamma\left(\zeta_v^{-2} \mid a_\zeta, b_\zeta\right)$ | Hyperprior: inverse-gamma distribution on $\zeta_v$ - defined per voxel, with hyperparameters<br>$a_\zeta$ and $b_\zeta$<br>See subject-level table for details of $\eta_m^{(s)}$ |
| | Probability that a given weight is drawn from signal rather than noise distribution | $\pi_{vm}$<br>$p(\pi_{vm}) = \beta(\pi_{vm} \mid a_\pi, b_\pi)$ | Hyperprior: Beta distribution with hyperparameters<br>$a_\pi$ and $b_\pi$ |
| Group Temporal Model | Temporal precision matrix to encourage consistency across subjects/runs | $\beta$<br>$p(\boldsymbol{\beta}) = \mathscr{W}(\boldsymbol{\beta} \mid a_\beta, \boldsymbol{B}_\beta)$ | Hyperprior: Wishart distribution with hyper parameters $a_\beta$ and $B_\beta$ |
| Group Amplitude Model | Mean of amplitudes | $\mu_h$<br>$p(\boldsymbol{\mu_h}) = \prod_{m=1}^{M} \mathcal{N}\left((\boldsymbol{\mu_h})_m \mid \tau_{\boldsymbol{\mu_h}}, \gamma^2_{\boldsymbol{\mu_h}}\right)$ | Hyperprior: Gaussian distribution with hyperparameters $\square_{\mu_h}$ and $\gamma^2_{\mu_h}$ |
| | Covariance of amplitudes | $\Sigma_h$<br>$p(\boldsymbol{\Sigma_h}) = \mathscr{W}\left(\boldsymbol{\Sigma_h}^{-1} \mid a_h, \boldsymbol{B_h}\right)$ | Hyperprior: Wishart distribution with hyperparameters $a_h$ and $B_h$ |
| Noise Model | Noise does not need group level modeling. This refers to residuals in matrix factorisation done at subject-level only:<br>$p\left(\boldsymbol{D}^{(sr)} - \boldsymbol{P}^{(s)} \boldsymbol{H}^{(sr)} \boldsymbol{A}^{(sr)}\right)$ | | |

#### Table S2: PROFUMO model details at the subject level

| | | | |
|---|---|---|---|
| Matrix Factorization | $\boldsymbol{D}^{sr} = \boldsymbol{P}^{s}\boldsymbol{H}^{sr}\boldsymbol{A}^{sr} + \boldsymbol{\varepsilon}^{sr}$ $\quad \boldsymbol{D}^{sr} \in \mathbb{R}^{N_v \times N_t}, \boldsymbol{P}^{s} \in \mathbb{R}^{N_v \times N_m}, \boldsymbol{H}^{sr} \in \mathbb{R}^{N_m \times N_m}, \boldsymbol{A}^{sr} \in \mathbb{R}^{N_m \times N_t}$ | | |
| Subject Spatial Model | Subject signal component<br>(first component of a Double Gaussian Mixture Model) | $P^{(s)}_{vm}\,(\lvert q^{(s)}_{vm} = 1)$<br>$p\left(\boldsymbol{P}^{(s)}_{vm}\middle\vert q^{(s)}_{vm}=1\right) = \mathcal{N}\left(\boldsymbol{P}^{(s)}_{vm}\middle\vert \mu_{vm}, \sigma^2_{vm}\right)$ | Hyperprior: Gaussian distribution with hyperparameters $\mu_{vm}$ and $\sigma_{vm}$ fed from the group model |
| | Subject noise component<br>(second component of a Double Gaussian Mixture Model) | $P^{(s)}_{vm}\,(\lvert q^{(s)}_{vm} = 0)$<br>$p\left(\boldsymbol{P}^{(s)}_{vm}\middle\vert q^{(s)}_{vm}=0\right) = \mathcal{N}\left(\boldsymbol{P}^{(s)}_{vm}\middle\vert 0, \left(\eta^{(s)}_m\right)^2\zeta^2_v\right)$ | Hyperprior: Zero-mean Gaussian distribution with variance hyperparameters<br>$\zeta_v$ fed from the group model; $\eta^{(s)}_m$ described below. |
| | Binary variable indicating whether a voxel belongs to the signal vs noise component. | $q^{(s)}_{vm}$<br>$p\left(q^{(s)}_{vm}\right) = \left(\pi_{vm}\right)^{q^{(s)}_{vm}}\left(1-\pi_{vm}\right)^{1-q^{(s)}_{vm}}$ | Hyperparameter $\pi_{vm}$ fed from group membership probability |
| | Extra parameter to capture variations in noise levels across voxels and modes | $\eta^{(s)}_m$<br>$p\left(\eta^{(s)}_m\right) = \mathcal{N}\left(\eta^{(s)}_m\middle\vert 0, \gamma^2_\eta\right)$ | Hyperprior: zero-mean Gaussian distribution with variance $\gamma^2_\eta$ |
| Subject Temporal Model | Signal Timecourse (per mode mode per subject per run)<br>Signal timecourse modeling is informed be HRF-induced autocorrelation structure (K) and between-mode precision matrix $\alpha$ | $A^{(sr)}_m$<br>$p\left(\boldsymbol{A}^{(sr)}_m\middle\vert\alpha\right) = \mathcal{N}\left(\boldsymbol{A}^{(sr)}_m\middle\vert\boldsymbol{0}, \alpha^{-1}\boldsymbol{K}_{\boldsymbol{A}}\right)$ | Hyperprior: Gaussian distribution, with mean 0 and variance $\alpha^{-1}K_A$ |
| | Noise Timecourse (per mode mode per subject per run per timepoint) | $\xi^{(sr)}_{mt}$<br>$p\left(\boldsymbol{\xi}^{(sr)}_{mt}\middle\vert\omega^{(sr)}_m\right) = \mathcal{N}\left(\boldsymbol{\xi}^{(sr)}_{mt}\middle\vert 0, {\omega^{(sr)}_m}^{-1}\right)$<br>$p\left(\omega^{(sr)}_m\right) = \Gamma\left(\omega^{(sr)}_m\middle\vert a_\omega, b_\omega\right)$ | Drawn from zero-mean, white Gaussian noise. Gamma hyperprior on noise precision ($\omega$) with hyperparameters $a_\omega$ and $b_\omega$ |
| | Temporal precision matrix (per subject, can also be defined per run). This combines the HRF-induced autocorrelation structure ($K_B$) with a prior on the between-mode precision matrix $\alpha^{(sr)}$<br>See Harrison et al., 2015 for details of a canonical double-gamma HRF modeling in PROFUMO. | $B^{(sr)}$<br>$p\left(\boldsymbol{B}^{(sr)}\middle\vert\boldsymbol{\alpha}^{(sr)}\right) = \mathcal{MN}\left(\boldsymbol{B}^{(sr)}\middle\vert\boldsymbol{0}, {\boldsymbol{\alpha}^{(sr)}}^{-1}, \boldsymbol{K}_B\right)$<br>$p\left(\boldsymbol{\alpha}^{(sr)}\middle\vert\boldsymbol{\beta}\right) = \mathcal{W}\left(\boldsymbol{\alpha}^{(sr)}\middle\vert a_{\boldsymbol{\alpha}}, \boldsymbol{\beta}\right)$ | Hyperprior for B: multivariate normal distribution with hyperparameters 0, $\alpha^{(sr)}$ and $K_B$<br>Hyperprior for $\alpha^{(sr)}$: Wishart distribution with hyperparameters $a_\alpha$ and $\beta$, the latter is fed from the group model. |
| Subject Amplitude Model | Subject- and run-specific amplitudes | $H^{(sr)} = diag(h^{(sr)})$<br>$p\left(\boldsymbol{h}^{(sr)}_m\middle\vert\boldsymbol{\mu}_h, \boldsymbol{\Sigma}_h\right) = \mathcal{N}\left(\boldsymbol{h}^{(sr)}_m\middle\vert\boldsymbol{\mu}_h, \boldsymbol{\Sigma}_h\right)$ | Hyperprior: Gaussian distribution with hyperparameters $\mu_h$ and $\Sigma_h$ fed from the group |
| Subject Noise Model | Noise model to ensure that noise variance is the same in every voxel | $\epsilon^{(sr)}$<br>$p\left(\varepsilon^{(sr)}\right) = \mathcal{MN}\left(\varepsilon^{(sr)}\middle\vert\boldsymbol{0}, \left(\psi^{(sr)}\right)^{-1}\boldsymbol{I}_V, \boldsymbol{I}_T\right)$<br>$= p\left(\boldsymbol{D}^{(sr)} - \boldsymbol{P}^{(s)}\boldsymbol{H}^{(sr)}\boldsymbol{A}^{(sr)}\right)$<br>$p\left(\psi^{(sr)}\right) = \Gamma\left(\psi^{(sr)}\middle\vert a_\psi, b_\psi\right)$ | Zero mean, white Gaussian noise. Gamma hyperprior on noise precision with hyperparameters<br>$a_\psi$ and $b_\psi$ |

**Figure S1: Patterns of network overlap for all 20 network pairs**

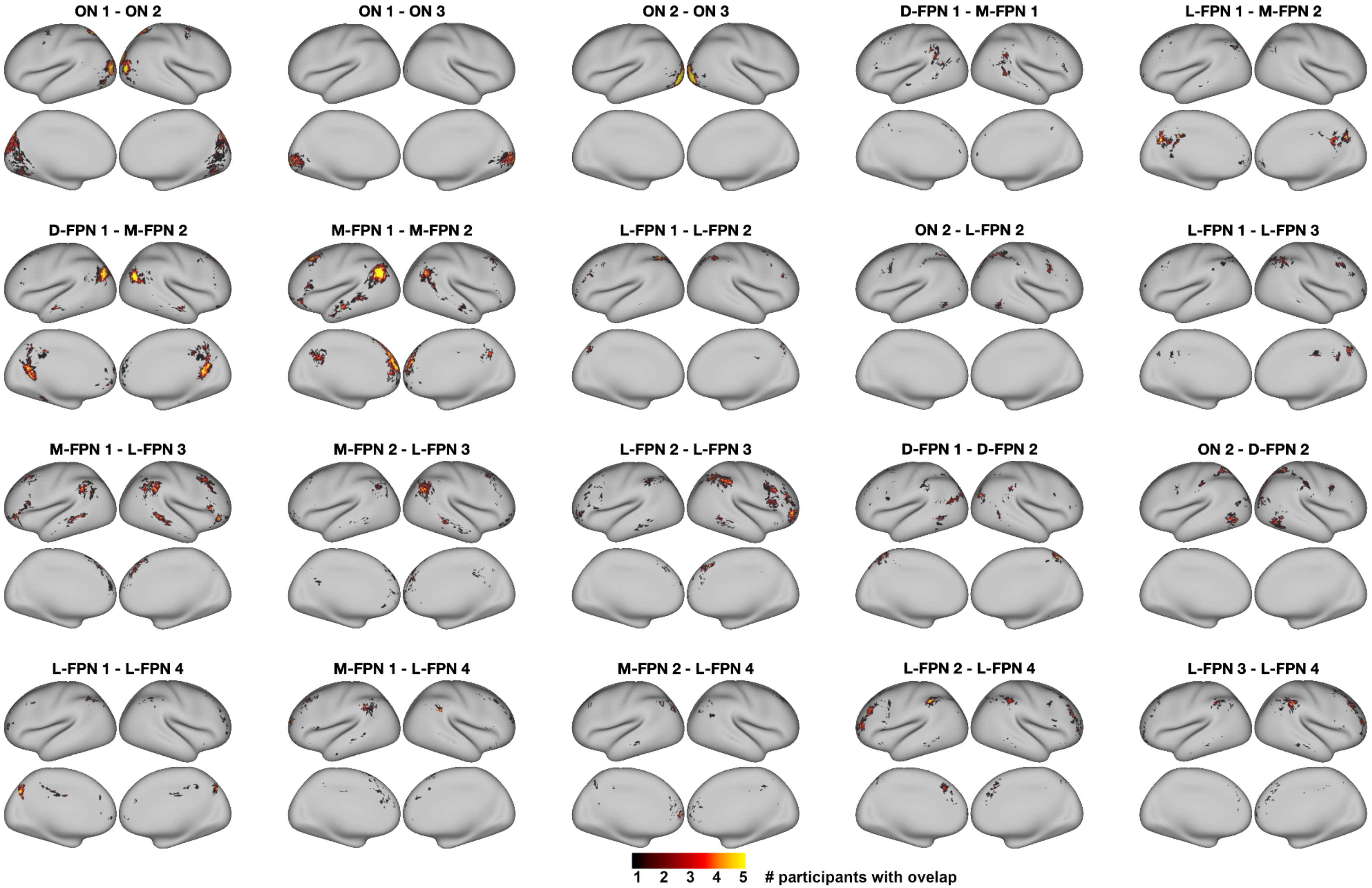


*Brain regions of spatial overlap between all 20 2-network pairs are displayed. At each vertex, the value represents the number of participants with overlap.*

**Figure S2: Interdigitation patterns for all 20 network pairs in example subject 195041**

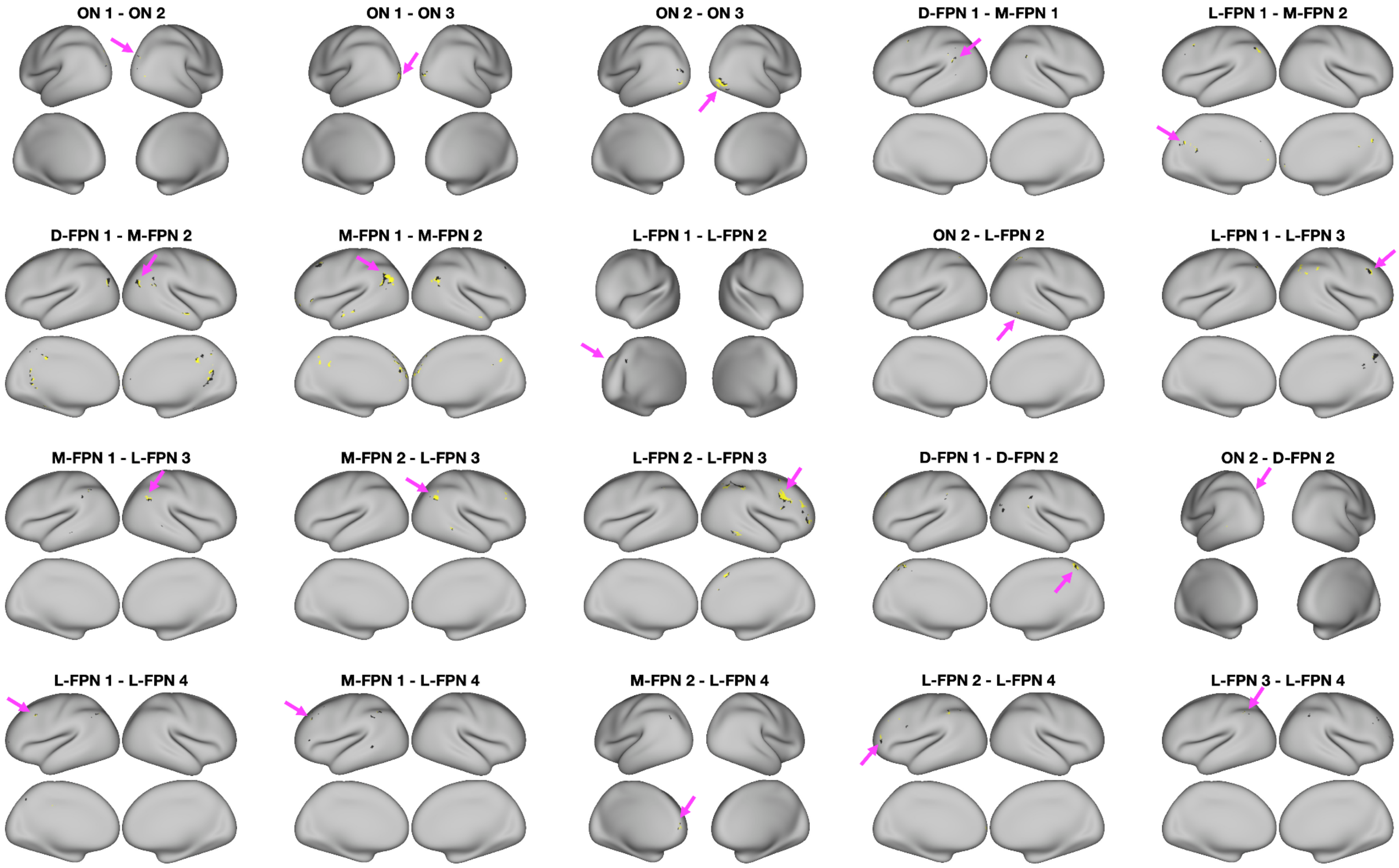


*Vertex assignments between network 1 (black) and network 2 (yellow) for all network pairs. Results for subject 195041 are shown as an example.*

**Figure S3: Mode stability across dimensionalities**

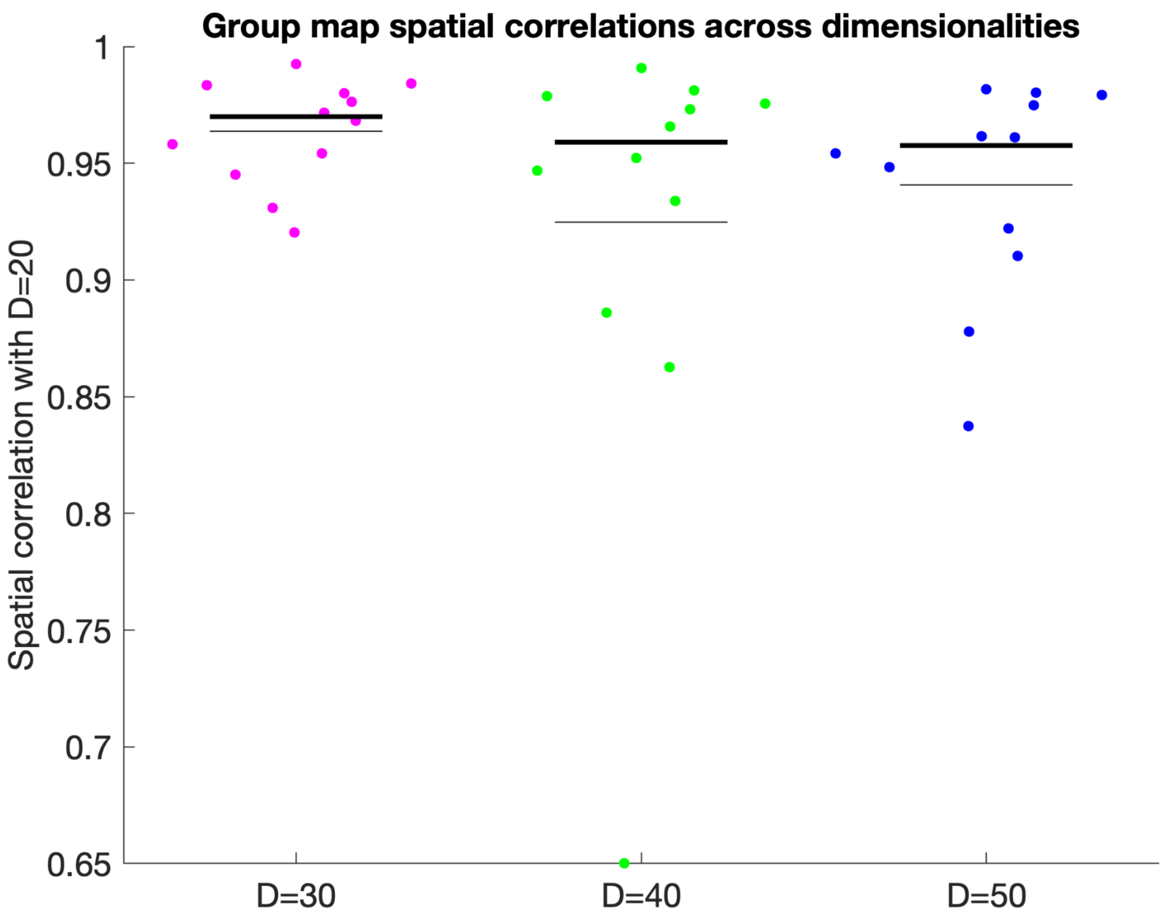


*The resting state networks investigated in this article were consistently found across a range of PFM dimensionalities.*

**Figure S4: Missing modes**

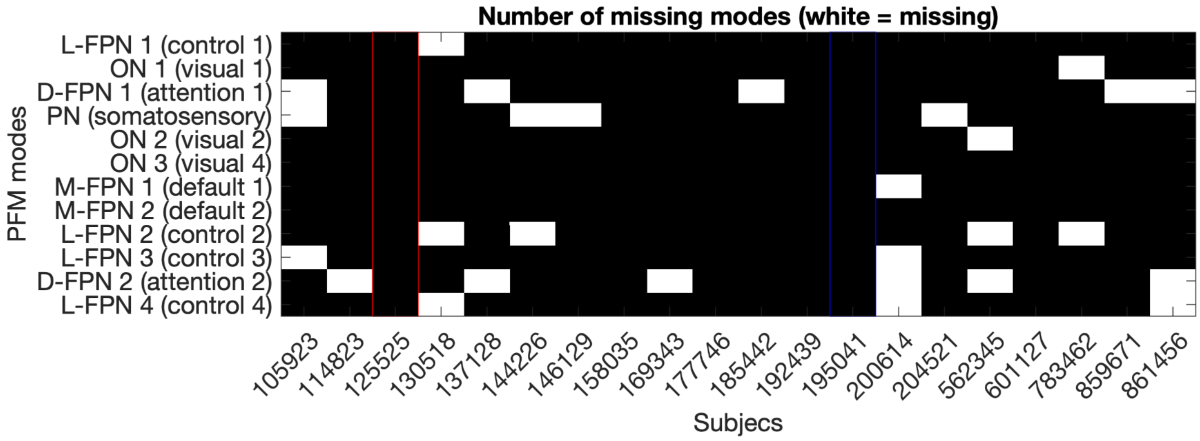


*Missing (non-estimable) modes for each subject based on the individual subject PROFUMO runs are shown in white. Red and blue columns indicate the examples participants used throughout this paper.*

**Figure S5: Subject maps**

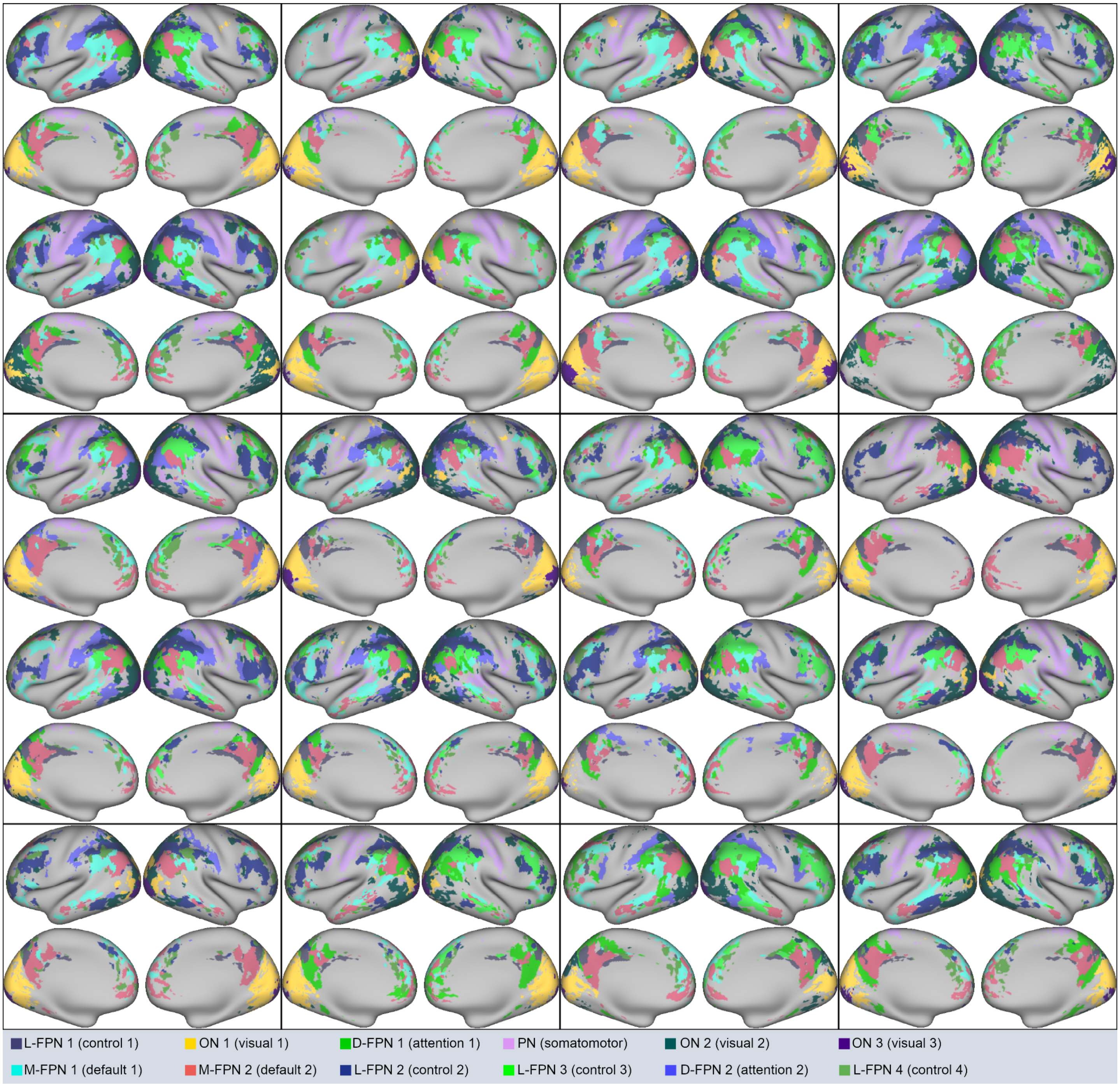


*Subject maps from individual subject PROFUMO runs. Vertices were allocated to the strongest network (i.e., overlap is not shown; see Fig. S5 for overlap maps). Vertices with a weight lower than one for each of the twelve modes were unassigned to focus on robust networks. Twin pairs are grouped vertically within each rectangle for the top rows and non-twin individual participants are shown on the bottom row. For naming purposes, modes were spatially mapped onto the Yeo-7 parcellation (Yeo et al., 2011) and we followed the naming convention from (Uddin et al., 2019). FPN = frontoparietal network; L = lateral; D = dorsal; M = medial; ON = occipital network; PN = pericentral network.*

**Figure S6: Unthresholded version of spatial similarity figure**

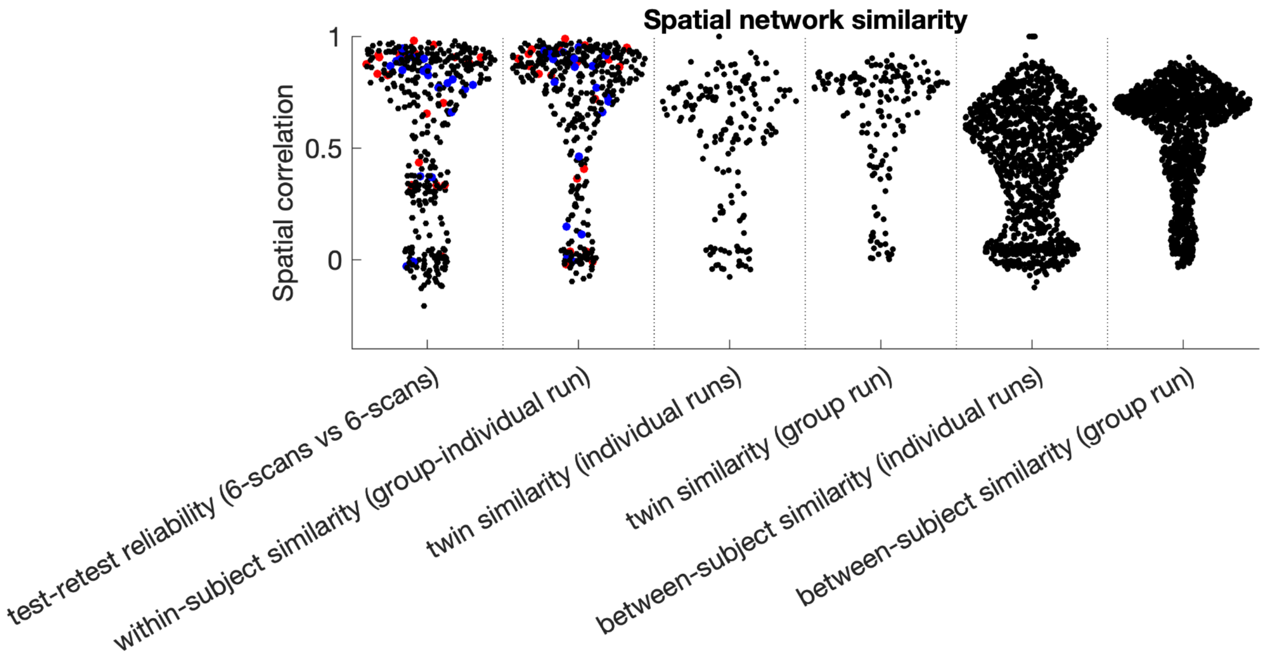


*Version of figure 5 in the main manuscript without removing any PROFUMO modes (either as missing modes at the individual participant level or as non-replicable modes at the group level). As expected, the tails of the distributions are heavier than observed in Fig. 5, but the column-wise differences remain evident.*

**Figure S7: Stability of temporal connectivity matrices**

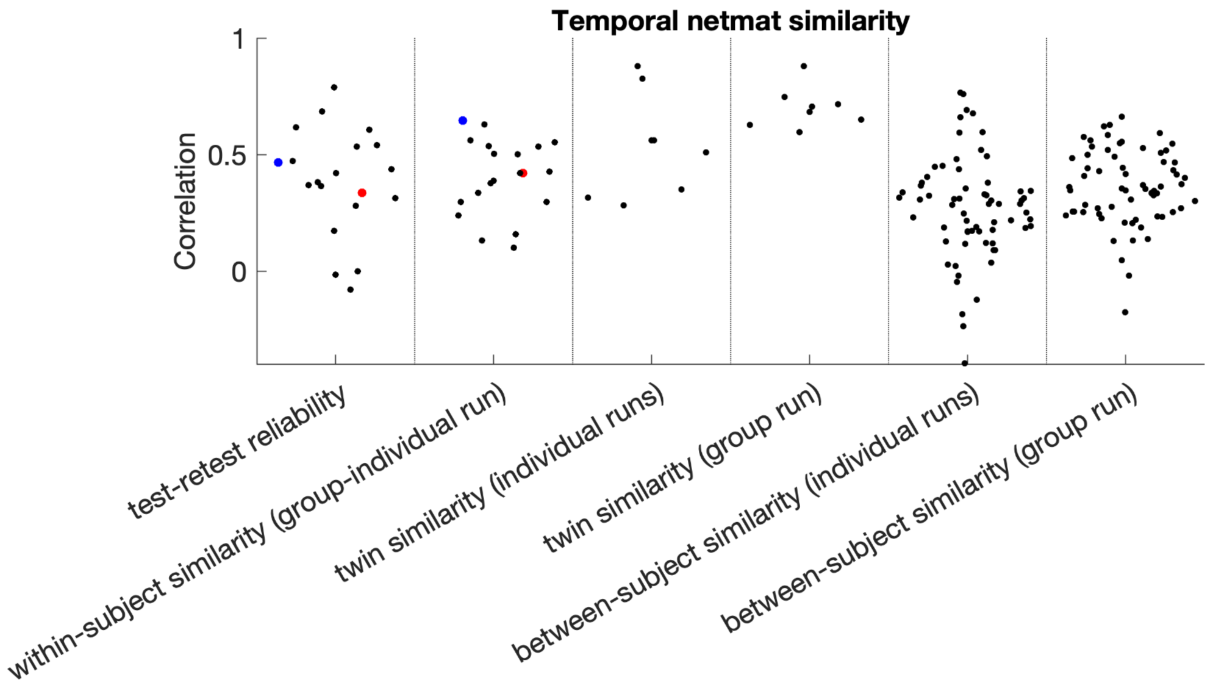


*Stability of temporal connectivity matrix (also known as network matrix or 'netmat'). Red and blue dots indicate results from the examples participants used throughout this paper.*

**Figure S8: Stability of spatial overlap matrices**

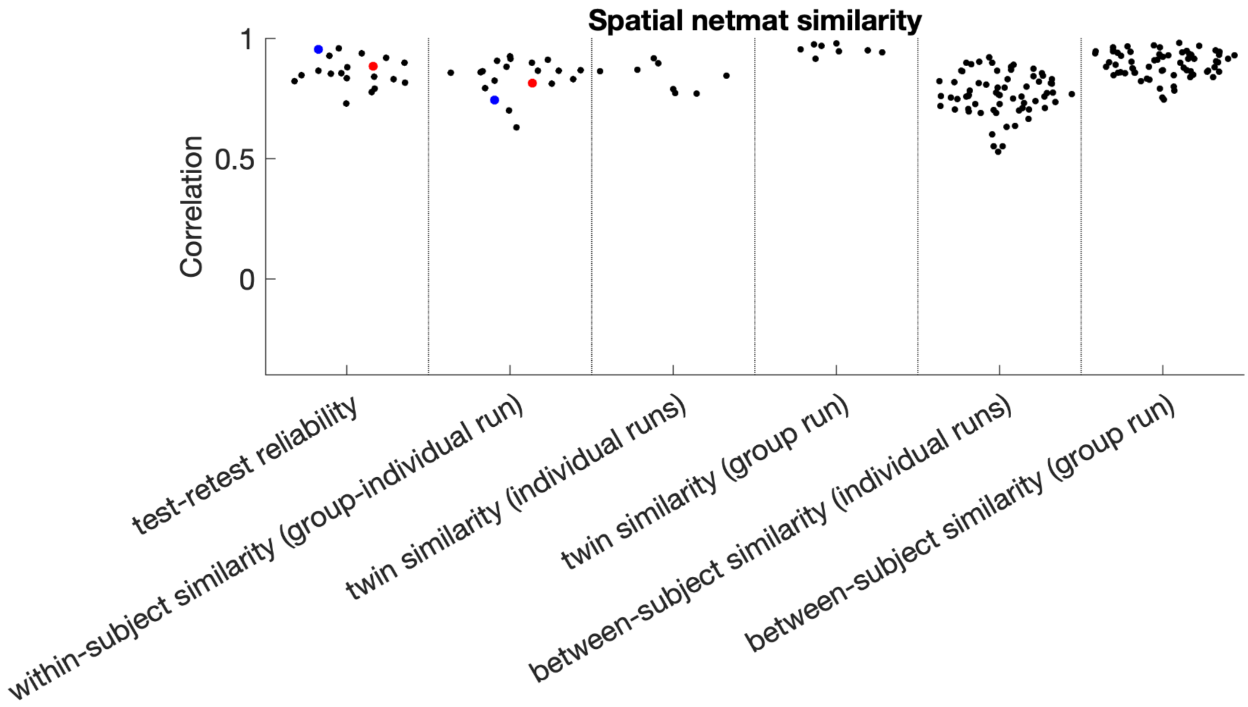


*Stability of spatial overlap matrix. Red and blue dots indicate results from the examples participants used throughout this paper.*

**Figure S9: Spatial overlap maps for all participants**

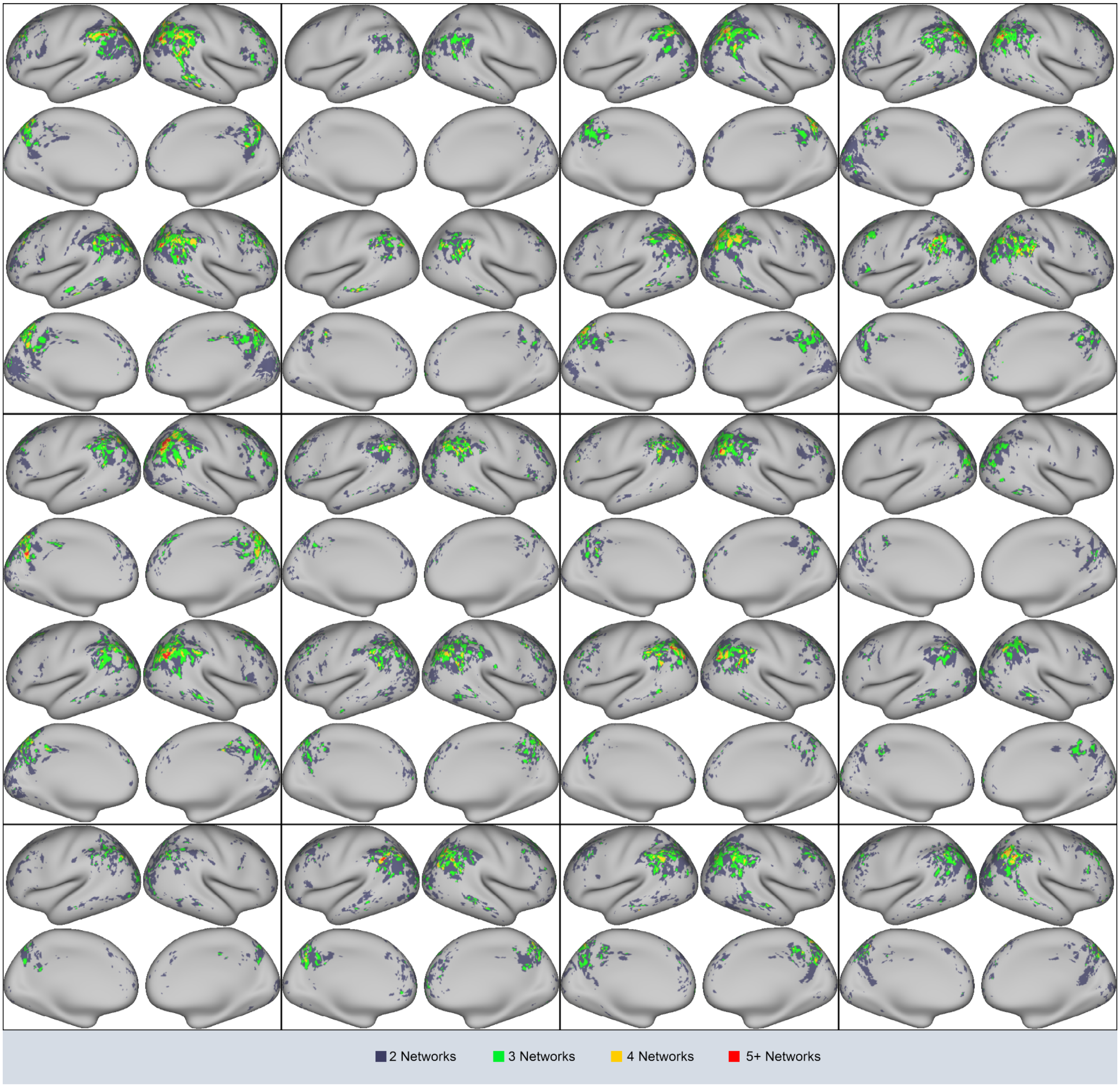


*Overlap maps for all participants. Twin pairs are grouped vertically within each rectangle for the top rows and non-twin individual participants are shown on the bottom row.*

**Figure S10: Timeseries similarity displayed separately for each network pair**

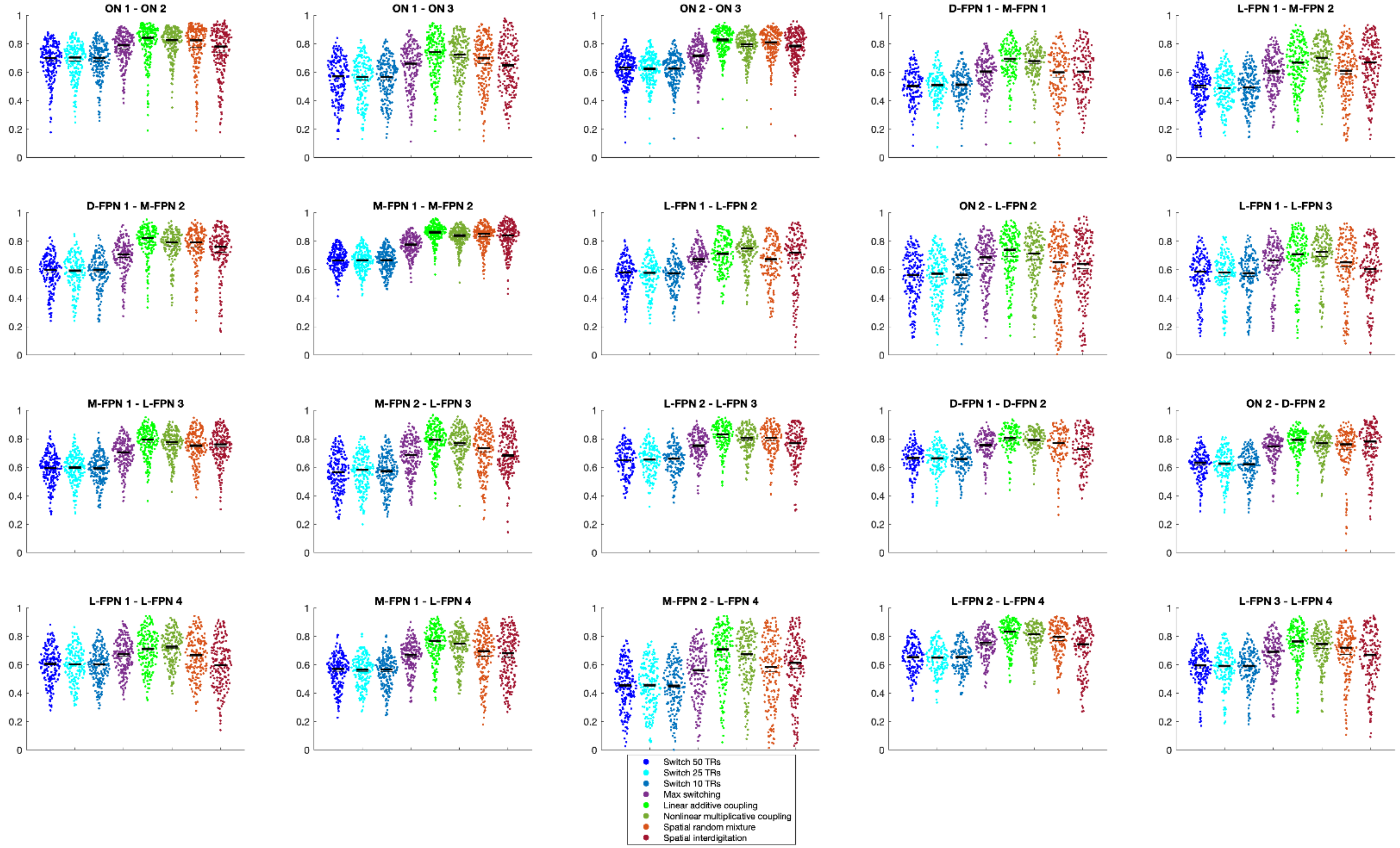


*Version of Figure 8 from the main manuscript with results shown separately for each of the 20 overlapping networkk pairs. Y-axis shows correlations between the true overlap timeseries and different versions of the simulated overlap timeseries. Thin black lines indicate the mean and thick black lines indicate the median. Highest similarity was consistently observed for the linear additive coupling simulated overlap timeseries in bright green.*

#### Figure S11: Frequency characteristics of semi-simulated timeseries

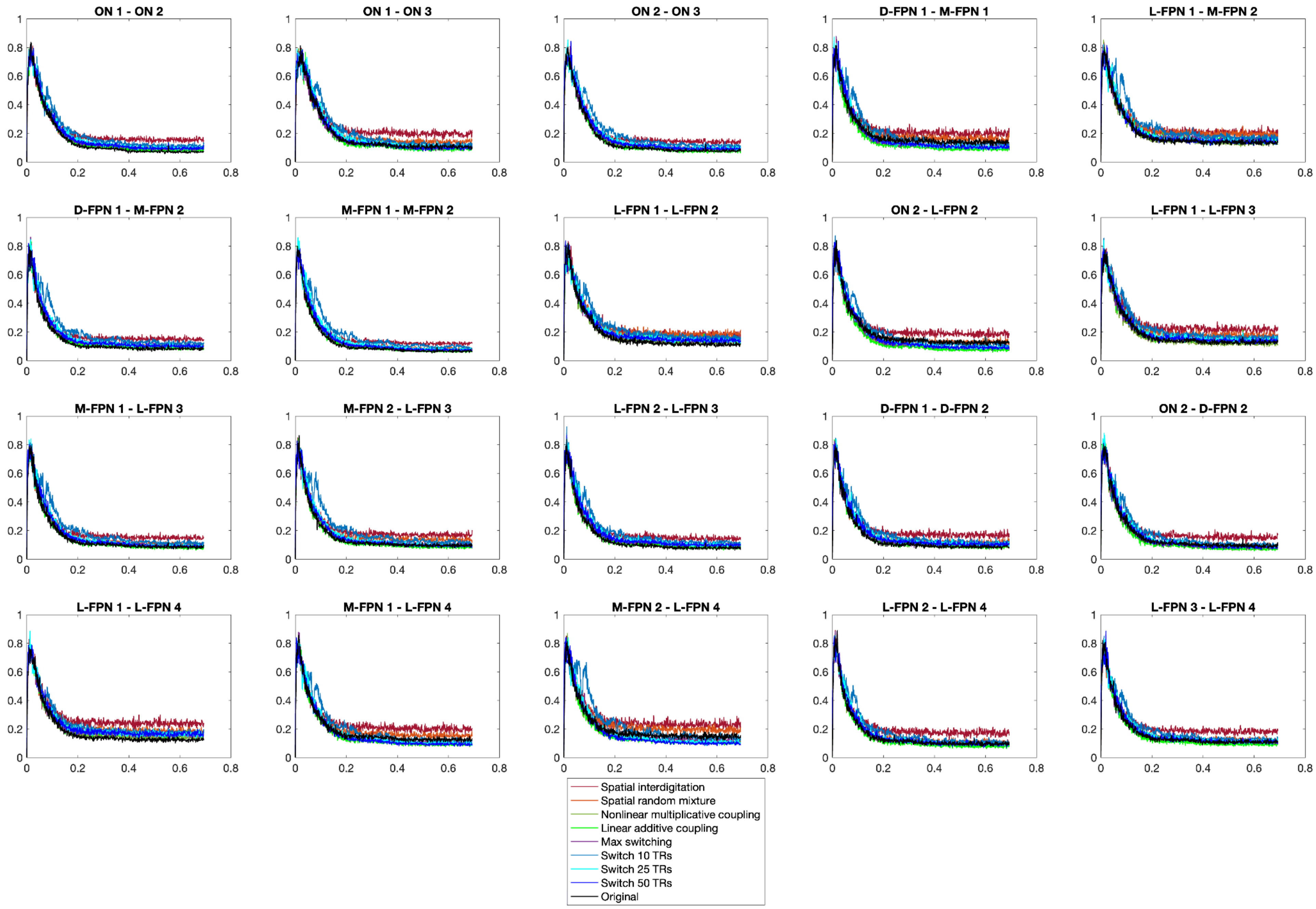

**Figure S12: HMM state summary**

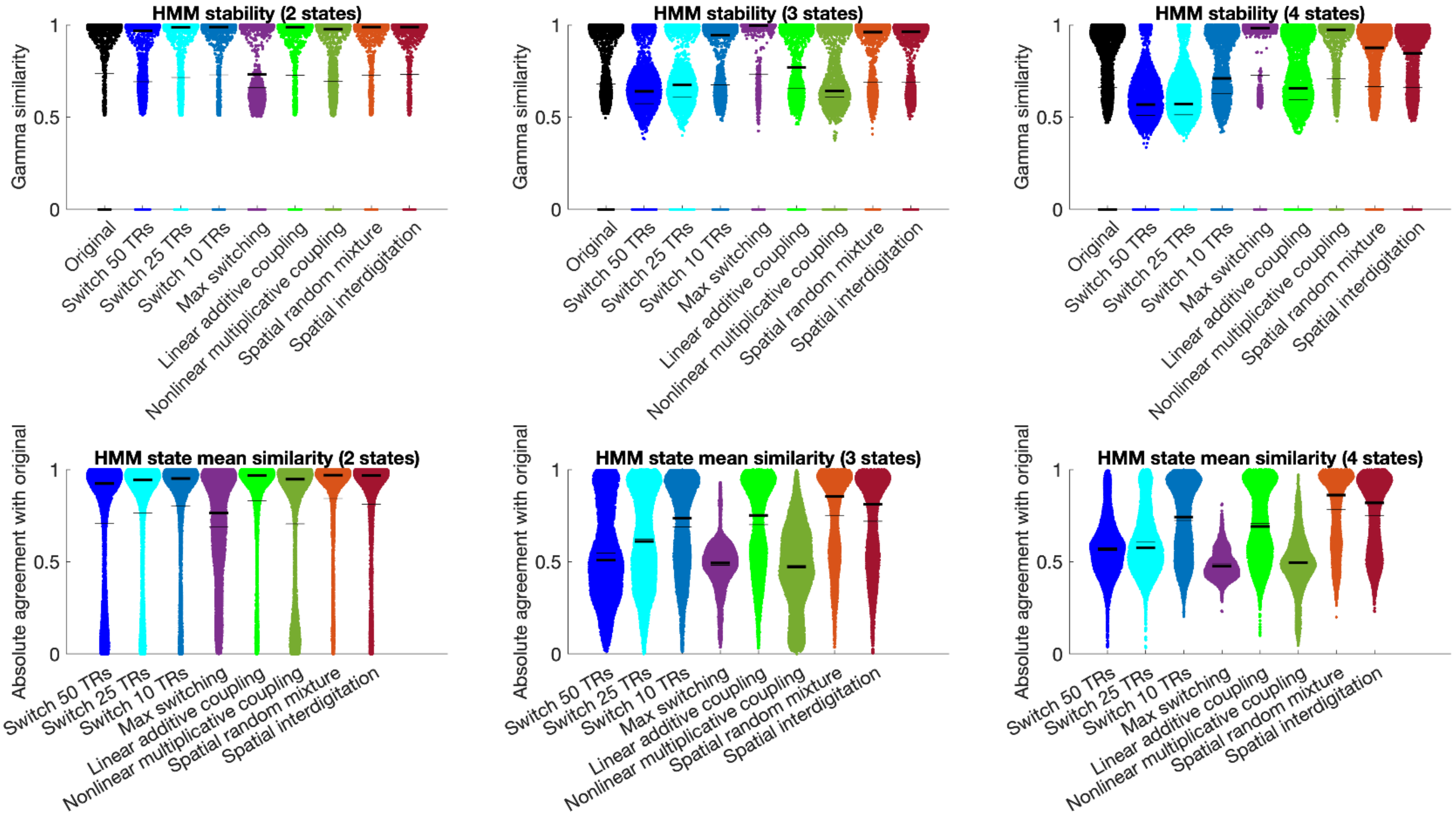


*The top row of figures displays the stability of HMM state estimation for the true overlap timeseries (black) and each of the different versions of the simulated overlap timeseries. The second row of figures displays the absolute agreement of HMM means between semi-simulated overlap timeseries and the original overlap timeseries after state reordering. Column reflect the number of states (2, 3, 4). Thin black lines indicate the mean and thick black lines indicate the median. Data were combined across all network pairs and all participants.*

#### Figure S13: Similarity of HMM state means between simulated and original overlap timeseries (2 states)

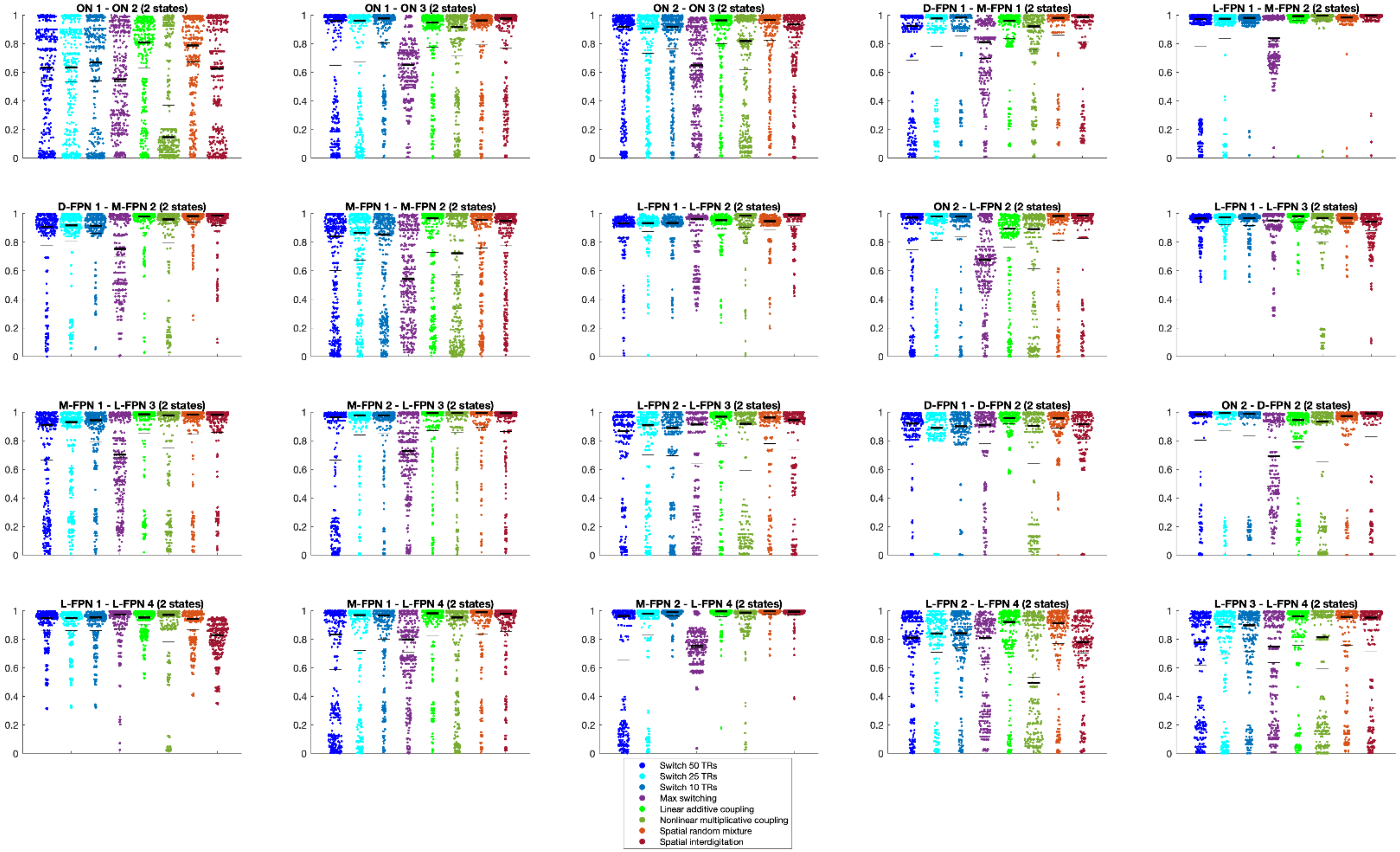


*Absolute agreement between HMM state means using the true overlap timeseries compared with different versions of the simulated overlap timeseries. Results for the 2-state solutions are shown separately for each network pair. Thin black lines indicate the mean and thick black lines indicate the median.*

**Figure S14: Similarity of HMM state means between simulated and original overlap timeseries (3 states)**

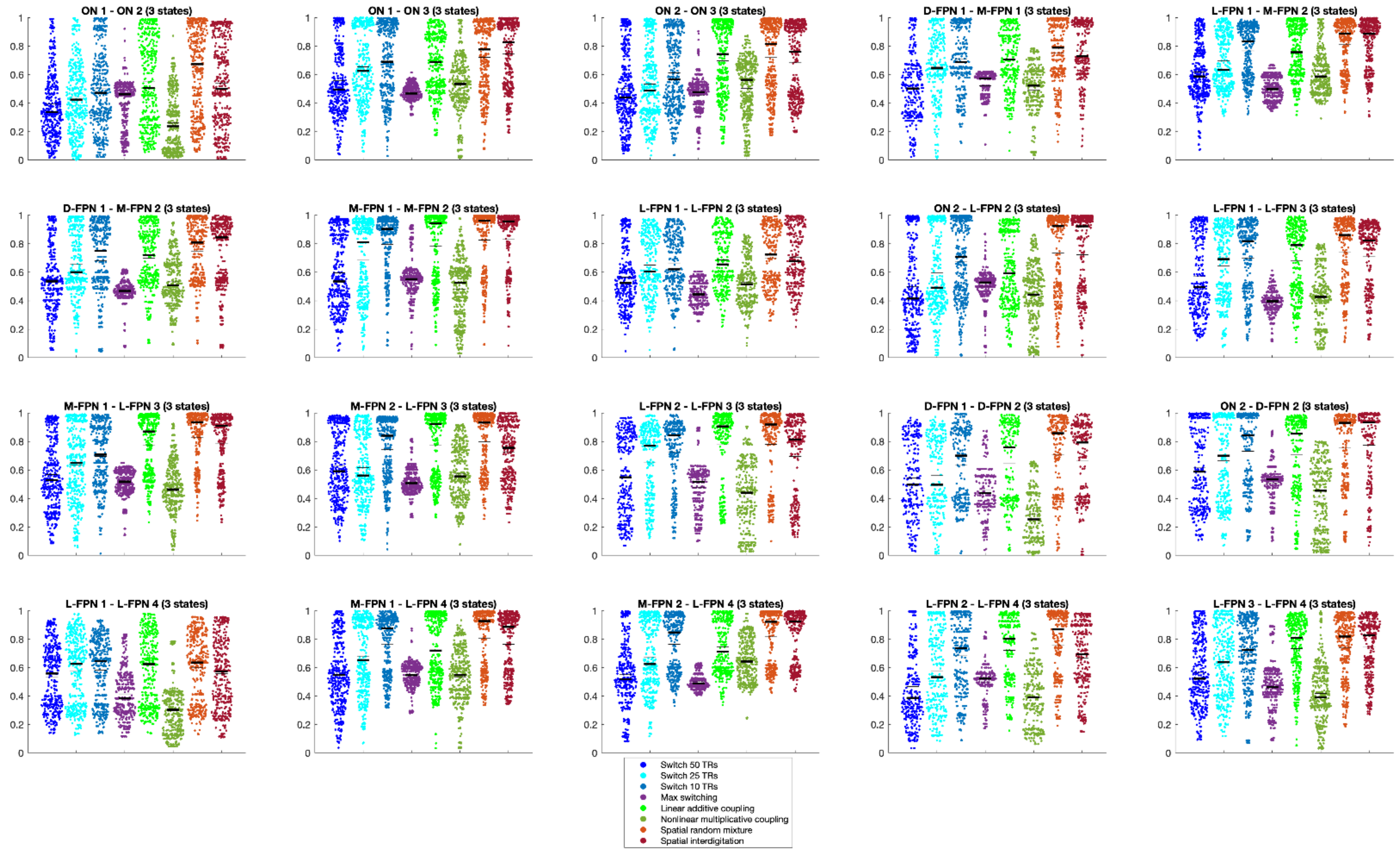


*Absolute agreement between HMM state means using the true overlap timeseries compared with different versions of the simulated overlap timeseries. Results for the 3-state solutions are shown separately for each network pair. Thin black lines indicate the mean and thick black lines indicate the median.*

**Figure S15: Similarity of HMM state means between simulated and original overlap timeseries (4 states)**

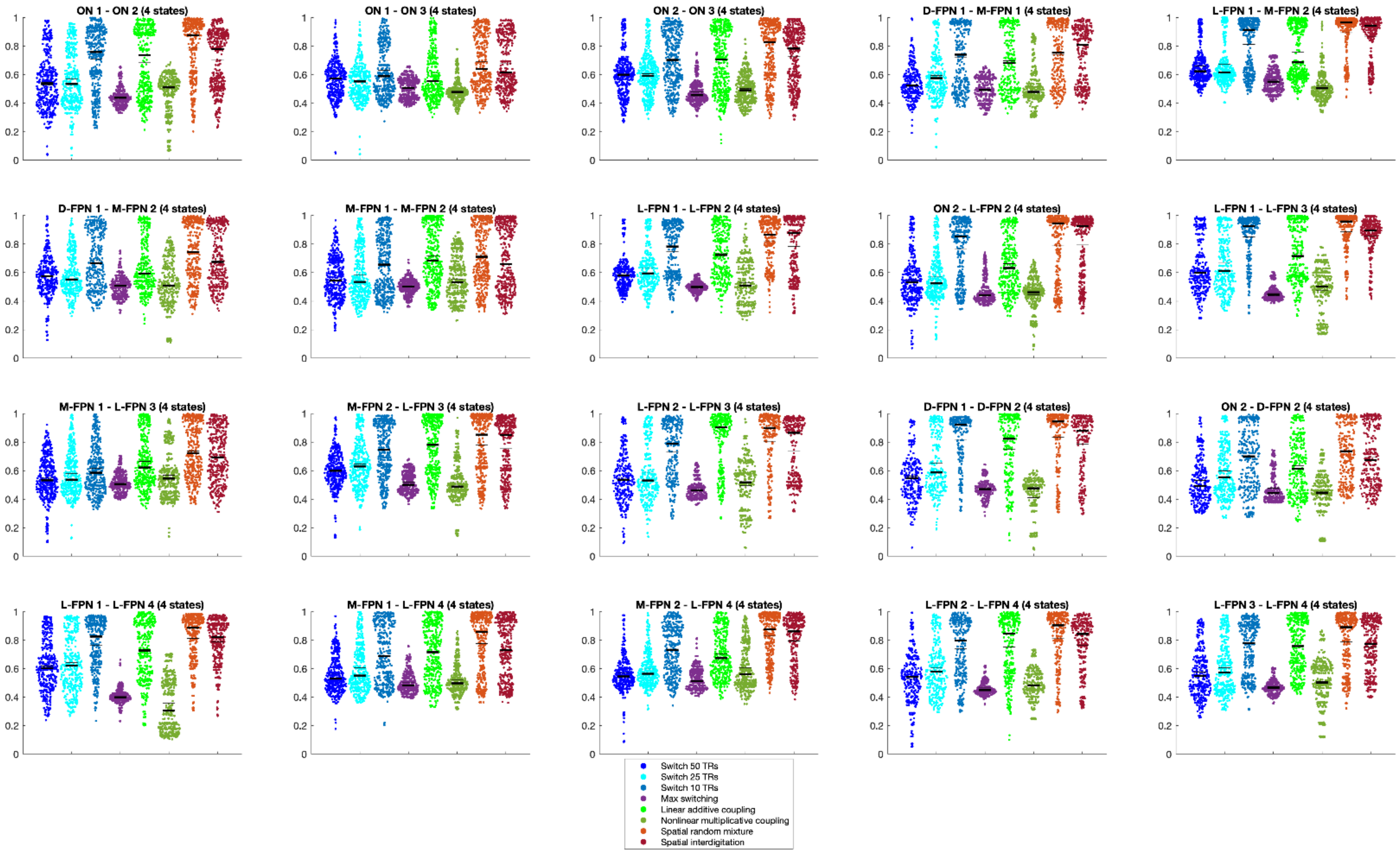


*Absolute agreement between HMM state means using the true overlap timeseries compared with different versions of the simulated overlap timeseries. Results for the 4-state solutions are shown separately for each network pair. Thin black lines indicate the mean and thick black lines indicate the median.*

#### References


Uddin, L.Q., Yeo, B.T.T., Spreng, R.N., 2019. Towards a Universal Taxonomy of Macro-scale Functional Human Brain Networks. Brain Topogr. https://doi.org/10.1007/s10548-019-00744-6

Yeo, B.T.T., Krienen, F.M., Sepulcre, J., Sabuncu, M.R., Lashkari, D., Hollinshead, M., Roffman, J.L., Smoller, J.W., Zöllei, L., Polimeni, J.R., Fischl, B., Liu, H., Buckner, R.L., 2011. The organization of the human cerebral cortex estimated by intrinsic functional connectivity. J. Neurophysiol. 106, 1125–1165.
